# Spatially resolved transcriptomics reveals plant host responses to pathogens

**DOI:** 10.1101/720086

**Authors:** Michael Giolai, Walter Verweij, Ashleigh Lister, Darren Heavens, Iain Macaulay, Matthew D. Clark

## Abstract

**Background:** Thorough understanding of complex model systems requires the characterisation of processes in different cell types of an organism. This can be achieved with high-throughput spatial transcriptomics at a large scale. However, for plant model systems this is still challenging as suitable transcriptomics methods are sparsely available. Here we present Grid-seq, an easy to adopt, micro-scale spatial-transcriptomics workflow that allows to study expression profiles across small areas of plant tissue at a fraction of the cost of existing sequencing-based methods.

**Results:** We compare the Grid-seq method with widely used library preparation methods (Illumina TruSeq). In spatial experiments we show that the Grid-seq method is sensitive enough to identify expression differences across a plant organ. We further assess the spatial transcriptome response of *Arabidopsis thaliana* leaves exposed to the bacterial molecule flagellin-22.

**Conclusion:** We show that our method can be used to identify known, rapidly flagellin-22 elicited genes, plant immune response pathways to bacterial attack and spatial expression patterns of genes associated with these pathways.

## Background

Most model plants and all crops species are multicellular, consisting of multiple organs and cell types with a multitude of physiological states [1]. A thorough understanding of these complex systems requires the ability to dissect and characterise processes in the different organs and cell types. This is challenging, though recently multi-omics single-cell studies have been flourishing [2], but high-throughput, high-resolution methodologies that assess molecular conditions with spatial resolution are sparsely available [3–6].

Although some spatial and low-input transcriptome profiling methods have been developed for animal model organisms [3–5,7], these methods are difficult to transfer to plants [6,8]. In comparison to animal cells, plant tissues hold a series of additional challenges: the robust plant cell wall requires specialised sample preparation (which makes reproducible, high-throughput sample preparation more difficult) and some plant secondary metabolites e.g. polyphenols can inhibit downstream enzymatic processes [9].For plants, single plant cells (protoplasts) can be obtained by enzymatic removal of plant-cell walls and subsequent fluorescent activated cell sorting (FACS) assays [10]. At the sub-cellular scale plant nuclei can be isolated within minutes by cell lysis and FACS [10–12]. However, ‘stimulus and response’ assays, such as differential gene-expression experiments or the characterisation of cell-type transcripts could be affected by these additional experimental procedures before RNA-extraction. Another important factor is the loss of spatial information when nuclei or protoplasts are extracted from a tissue. Thus methods such as fluorescent in-situ hybridisation (FISH) [13], laser-capture microdissection (LCM) [14–16] or the SPATiAL TRANSCRIPTOMICS [4,6] workflow are better suited to understand spatial transcription changes. However, all three methods need specific tissue preparations (e.g. cryo-sectioning, permeabilization or fixation) and specialised protocols to assess transcriptome levels: FISH methods require imaging of transcripts and are restricted to multiplexing a few fluorescent anti-sense probes at a time [3], LCM requires specialised equipment and training for precise, laborious excision of specific tissue elements [6] and the SPATiAL TRANSCRIPTOMICS protocol requires preparation of thinly sectioned, permeabilised samples and custom made DNA arrays [4,6].

Despite the high level of resolution that can be achieved with all these methods, they are not easily applied in most laboratories. We aimed to overcome this with our Grid-seq workflow. Grid-seq is designed to quickly process mechanically dissected samples into sequencing libraries using standard laboratory equipment and can be used in most modern laboratories. Grid-seq is based on three consecutive steps: (1) rapid, mechanical sample dissection of small e.g. 1 mm^2^ leaf areas, (2) a high-throughput method for high quality mRNA extraction of difficult to lyse plant tissues and (3) next generation sequencing (NGS) library construction (**Figure 1**).

**Figure 1.**
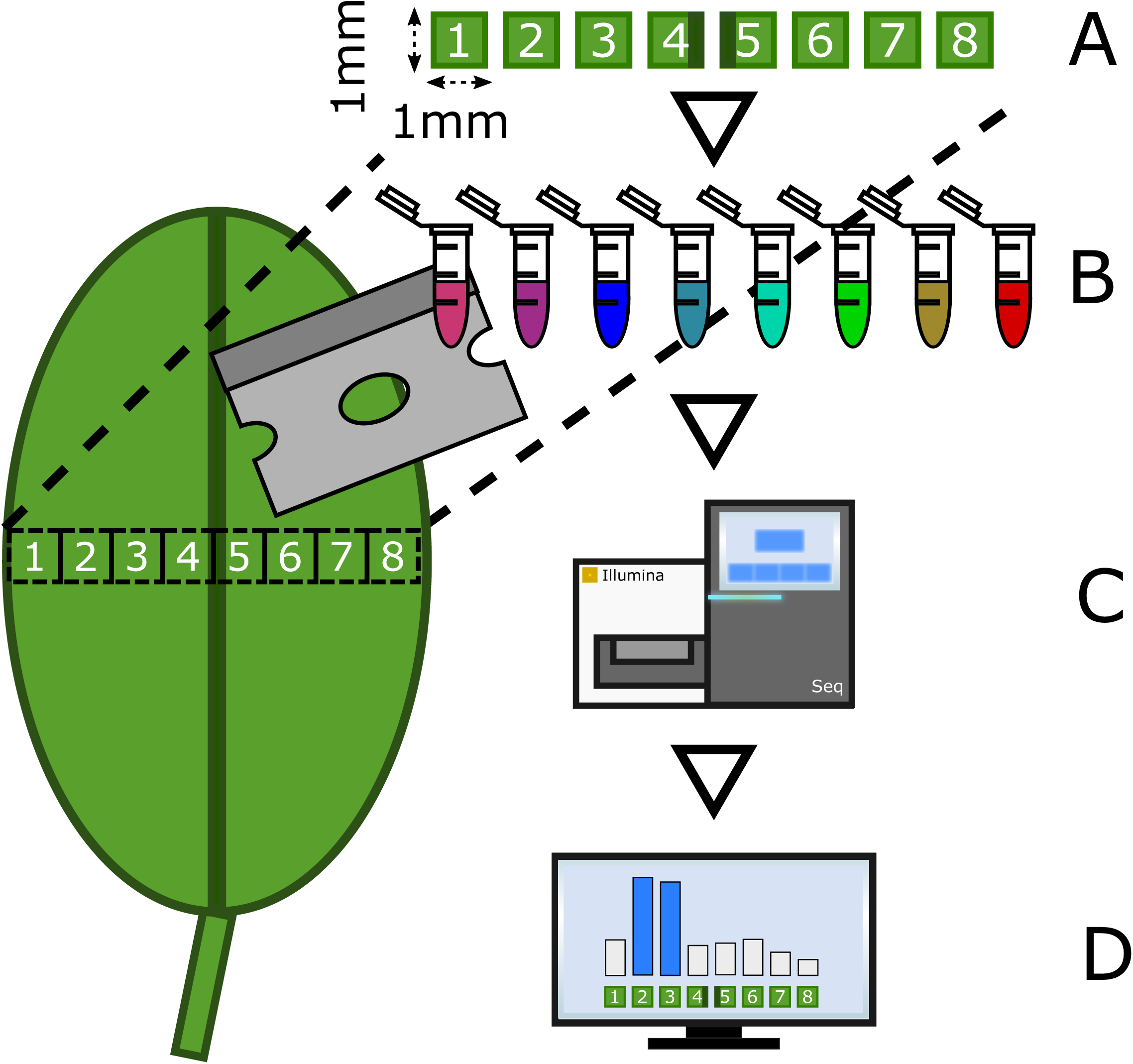
Overview of the Grid-seq workflow: (a) Tissue sections of approximately 1 mm^2^ size are mechanically extracted (e.g. a cross-section of a leaf) and after mRNA extraction (b) prepared into uniquely barcoded Illumina sequencing libraries. After (c) Illumina sequencing (d) transcript specific, spatial expression data can be assessed and analysed.

In a series of experiments, we compare the performance of Grid-seq with standard RNA-seq (Illumina TruSeq) experiments. Using Illumina sequencing we identify differentially expressed (DE) genes in 1D (across lateral leaf sections) observing how transcripts and expression levels vary across the tissues that make up the leaf organ. We compare large-scale vs. fine-scale transcriptome experiments ability to detect plant responses induced by the bacterial peptide flagellin-22 (flg22), a well-described pathogen-associated molecular pattern (PAMP) that triggers plant immune responses [17]. By comparing our data with published datasets for ‘<u>fla</u>gellin rapidly <u>e</u>licited’ (FLARE) genes [18] we identified 143 of 253 described FLARE genes that overlap with our data, and a further 428 genes with similar expression patterns to FLARE genes. We show that the detected 428 transcripts, are enriched for plant defence responses and that spatial transcriptome data can be used to reconstruct the spatial expression of pathway components across leaves. In additional pilot experiments using *Arabidopsis thaliana* and the oomycete pathogen *Albugo laibachii* we also show that the dual dataset of both host and pathogen, are captured, and spatial expression profiles can be constructed which could be used to study the biotic interactions in infections

## Results

### Does leaf dissection induce wounding response gene expression profiles?

Physical wounding of plants is known to induce wounding related gene expression [19]. This is an important point to consider as the Grid-seq workflow dissects tissue into ∼ 1 mm^2^ squares followed by immediate snap freezing on dry ice. Yet dissection could potentially lead to activation of wounding related gene expression and dissection takes longer as the resolution increases (grid size). As any wounding effect could form a technical limitation to Grid-seq we measured the number of DE-genes found after tissue dissection. For this we prepared ∼1 mm^2^ leaf squares (3 biological replicates per time-point) at the time-points: 0-minutes, 2.5-minutes, 5-minutes and 10-minutes between cutting and freezing on dry ice (when all enzymatic reactions cease). To determine the number of DE-genes at each time-point we compared the 2.5-minute, 5-minute and 10-minute samples with the 0-minute samples as an unwounded reference. This analysis showed just 1 DE-gene (AT2G37130) at the 2.5-minute time-point (which was not significant at later time-points) there were no DE-genes at the 5-minute time-point and 13 genes at the 10-minute time-point (see: **Additional_file1.docx** and **Additional_file2.xlsx**) suggesting that the transcriptional response to wounding starts between 5 and 10 minutes. We looked for enriched biological processes in the combined set of 14 genes and detected three genes at the 10-minute time-point being associated with the GO-term ‘response to wounding’: *TPS04, TAT3* and AT1G62660 (see: **Additional_file1.docx** and **Additional_file2.xlsx**). This indicates that it is highly desirable to cut and snap freeze sample material within 10 minutes to avoid perturbation of results – a time window in which we find that sample dissection is easily achievable.

### Spatially resolved transcriptomics data reveals leaf tissue specific gene expression

We assayed Grid-seq’s ability to detect known gene expression differences between tissue types in untreated leaves. Briefly, we dissected a lateral cross-section of an *A. thaliana* leaf (3 biological replicates) into a 1-dimensional (1D) expression map of eight circa1 mm^2^ squares (**Figure *2*A**). Each cross-section was sampled according to the same pattern: the leaf margins were located at square-1 and square-8 and the midvein at square-5. We then identified DE-genes by comparing the midvein with the lamina and the leaf margins with the lamina. This resulted in 393 DE-genes for the midvein and 686 DE-genes for the leaf margins comparison (**Figure 2** and file: **Additional_file2.xlsx**).

**Figure 2.**
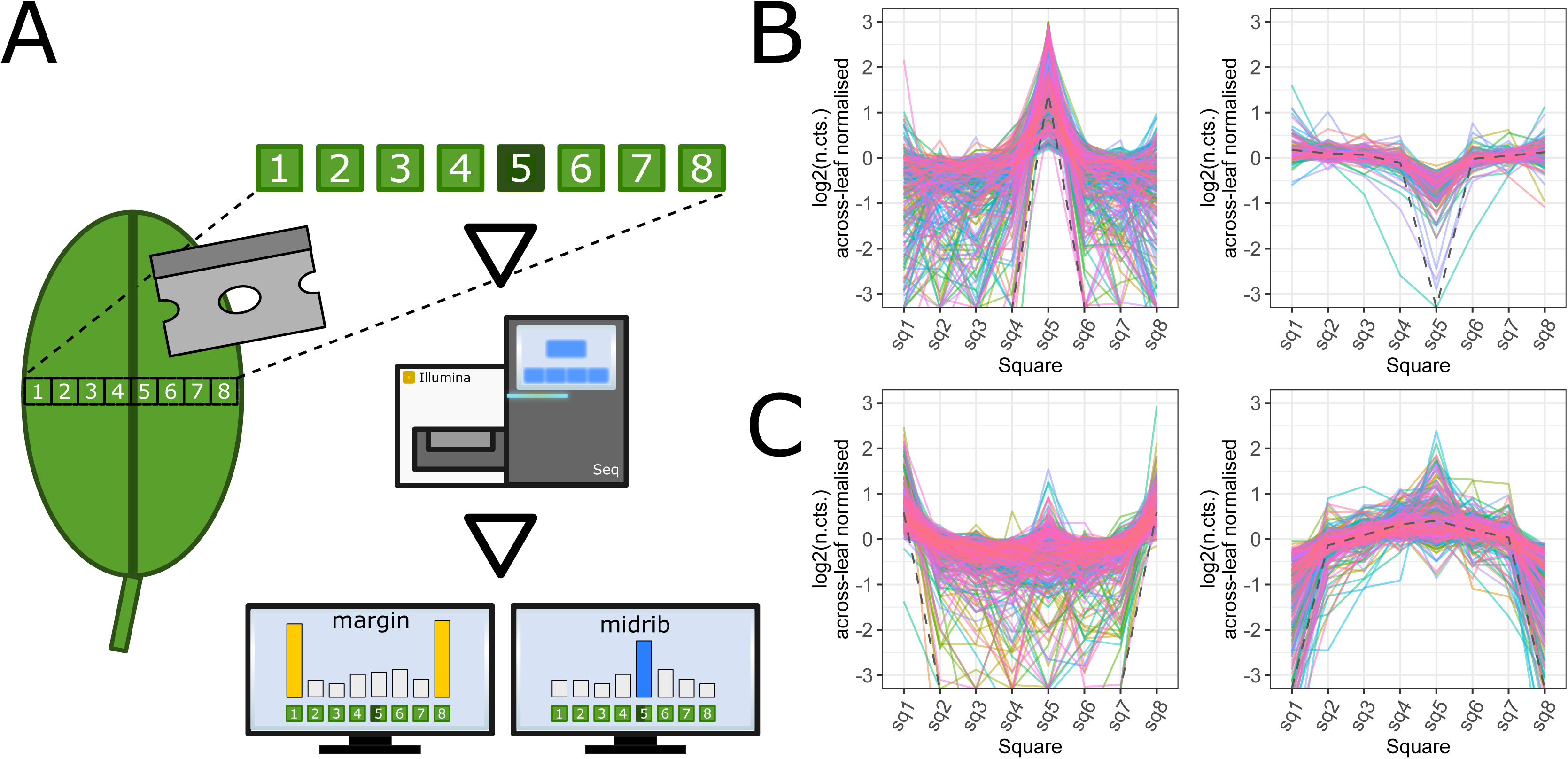
Identification of midvein and edge DE-genes in a lateral 1D leaf cross-section: (A) DE-gene analysis of a 1D *A. thaliana* leaf lateral cross sections by comparing the midvein (square-5) or the margin squares (square-1 and square-8) with the ‘bulk’ (remaining) leaf sections. The two images in (B) show 393 DE-genes with higher (left, 256 DE genes) or lower (right, 137 DE genes) expression values in the midvein. The images in (C) show 686 DE-genes higher (left, 403 DE genes) or lower (right, 283 DE genes) expression in the leaf margins. The grey dashed line in each plot (B and C) represents a trend-line for the average log2(counts) of all genes normalised across the leaf squares.

### Comparison of spatial and bulk transcriptomics after localised flg22 stimulation

To compare spatial (only treated areas) with bulk (large leaf areas with treated and untreated areas) PAMP immune responses we used a flg-22 syringe infiltration assay. For this we produced small, local infiltration spots on the abaxial, left-hand side of a leaf of 6 biological *A. thaliana* replicates using either 500 nM flg22 or water (**Figure 3A**). We incubated the plants for 1 hour and sampled by dissecting leaf samples with a 1D system as above, briefly: square-1 (in the middle of the left half of a leaf) as infiltration spot and then laterally towards the midvein square-2 as non-vascular leaf tissue, square-3 as the midvein and square-4 as non-vascular leaf tissue.

**Figure 3.**
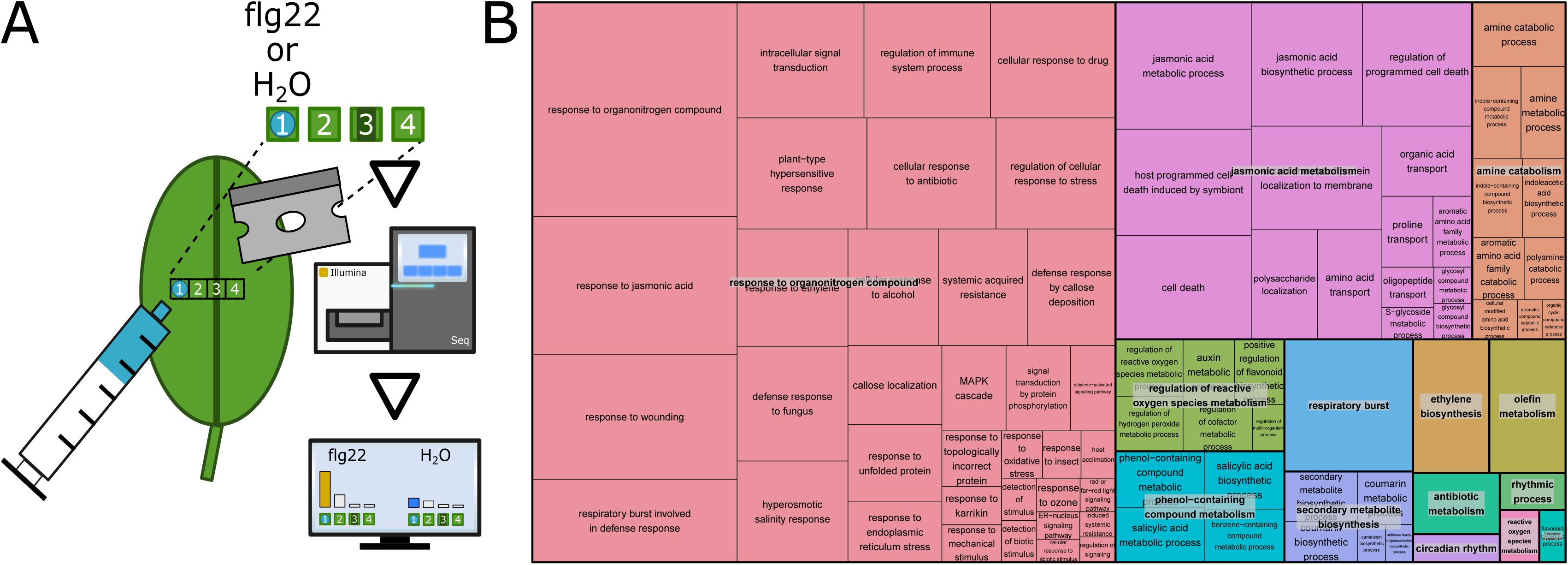
Elicitation of early *A. thaliana* defence response genes by infiltrating the bacterial peptide flg22: (A) To provide a strong stress stimulus we used syringe infiltration of either 500 nM flg22 or, as a control, water on a small area of the abaxial side of a leaf (square-1). 1 hour after infiltration we dissected 4 squares of a lateral leaf section with square-1 being the infiltration spot, square-2 and square-4 untreated, non-vascular leaf tissue and square-3 as midvein. The figure in (B) shows the REVIGO *[20]* treemap of the detected 168 GO-terms grouped under parent terms such as ‘response to organonitrogen compound’ (rose), ‘jasmonic acid metabolism’ (purple), ‘amine catabolism’ (orange) ‘regulation of reactive oxygen species metabolism’ (green), ‘phenol-containing compound metabolism’ (azure blue), ‘respiratory burst’ (blue), etc.. The size of each rectangle relates to the absolute log10(q-value) – the larger the more significant.

In our analysis we wanted to measure how bulk RNA-seq datasets compared to spatially collected ones by using the number of detectable DE-genes. We hypothesised that the spatial analysis of the small, local treatment spot (square-1) and its surroundings (square-2, square-3 and square-4) would reveal more and distinct types (or waves) of flg22 responsive DE-genes than a bulk analysis would – especially of rarer transcripts. To measure the effect of spatial information alone we simulated an *in silico* flg22 bulk experiment by combining the data from flg22 or water treated square-1 with the other untreated squares-2, 3 and 4 of the same leaf. We then called the treatment responsive DE-genes from the bulk files, detecting 65 DE-genes (39 higher expressed, 26 lower expressed) 1 hour after flg22 infiltration. We detected 887 more DE-genes (952 in total) by comparing the single squares of the flg22 and water infiltration dataset: 646 DE-genes for square-1 (416 higher, 230 lower expressed), 401 DE-genes for square-2 (306 higher, 95 lower expressed), 9 DE-genes for square-3 (8 higher, 1 lower expressed) and any DE-genes for square-4 (see: **Additional_file2.xlsx**). In contrast comparing the gene lists of the *in silico* bulk and spatial analysis we detected that 4 DE-genes were exclusively called from the bulk dataset and 64 genes were shared by both datasets.

To identify the biological processes uncovered by our transcriptomics experiments we performed a GO-term enrichment analysis the spatial flg22 related DE-gene datasets. From all (952) DE-genes we obtained 168 enriched GO-terms; among them we observed a high number of biological processes related to stress and defence responses (see: **Figure 3B**).

### Early elicited flg22 response genes of local, fine-scale stimulation

To simulate an initial pathogen encounter we used a milder stimulus method than the above described syringe infiltration: we prepared 6 biological *A. thaliana* replicates by depositing 1 µl of 500 nM flg22 on square-3 of the abaxial side of a leaf and 1 µl of water (internal control) on square-6 (equivalent locations due to leaf bilateral symmetry). After one hour we extracted the treated leaf area as a 1D lateral cross-section containing 8 separate 1 mm^2^ squares (**Figure 4A**). We were interested in DE-genes at the site of flg22 spotting (square-3) and in adjacent sections (square-2 and square-4) as we reasoned that the plant would respond to the PAMP locally at first and then responses via signalling to adjacent tissues and the rest of the plant. We called DE-genes by comparing the flg22 with the water droplet spots (square-3 vs square-6) and the adjacent sections with their corresponding bilateral equivalents (square-2 vs square-7 and square-4 vs square-5). Due to the milder stimulus in comparison to the flg22 infiltration dataset we expected the number of DE-genes could be lower than in the syringe infiltration experiment where we detected 952 DE genes. Indeed, we identified a lower number of 523 DE-genes (491 higher expressed, 32 lower expressed) for the droplet spot, and 5 DE-genes in the adjacent sections (1 higher expressed DE-gene in the square-4 square-5 comparison and 4 higher expressed DE-genes in the square-2 vs square 7 comparison). Thus, in total we detected 526 individual DE-genes.

**Figure 4.**
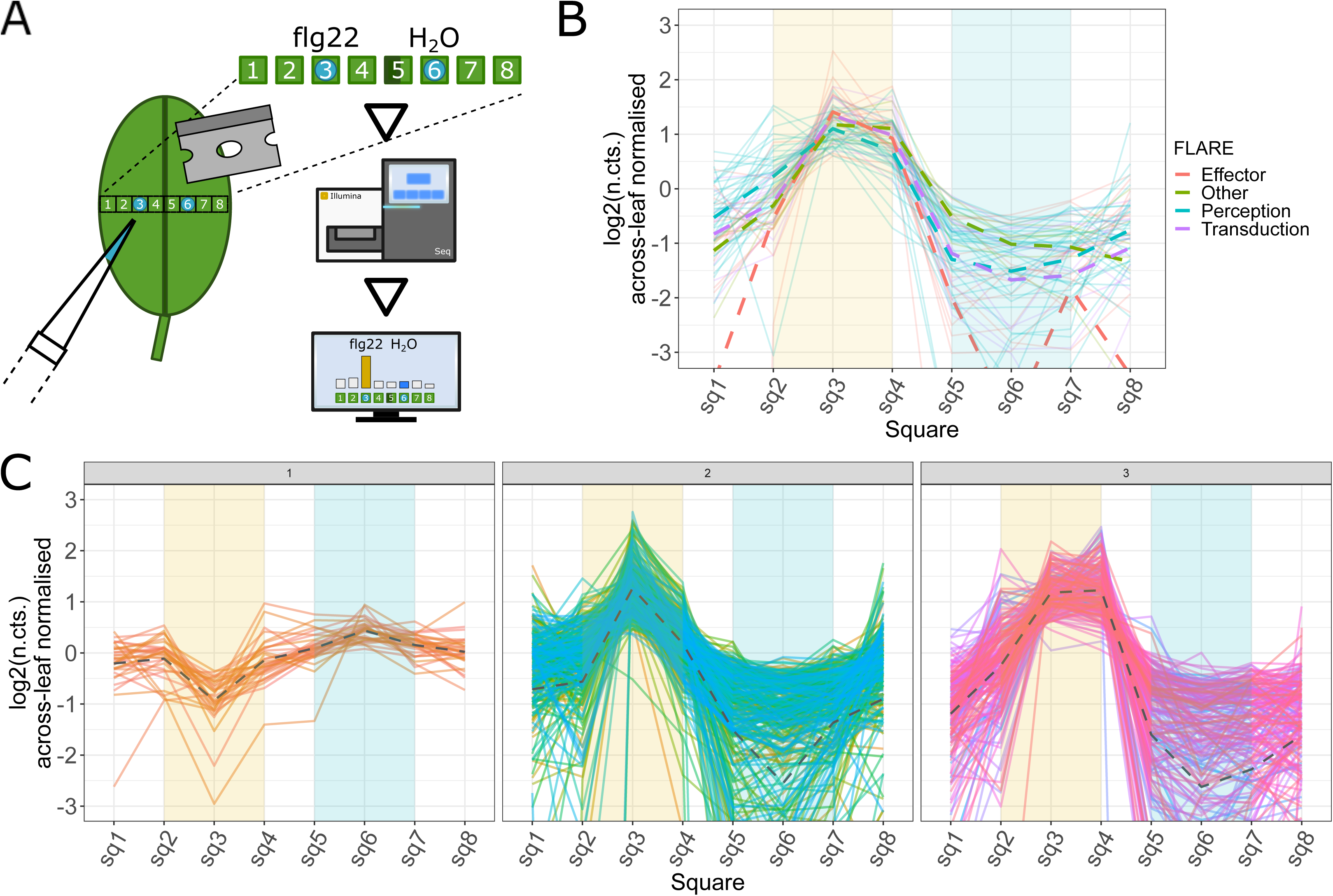
Elicitation of early *A. thaliana* defence response genes by droplet depositing the bacterial peptide flg22: (A) As a milder stress stimulus than flg22 syringe infiltration we pipetted a 1 µl droplet of 500 nM flg22 and, as an internal control, water on the abaxial surface of a leaf. 1 hour after droplet deposition we dissected a lateral section into 8 squares with square-1 and square-8 as leaf margins, square-3 as flg22 treated spot, square-6 as water treated spot and square-5 as midvein. Image (B) shows an overlay of the spatial expression patterns of the 63 FLARE genes characterised by Navarro et al. *[18]* present in our dataset. Each group is coloured separately, the average expression of each FLARE group is shown as the dashed line. Image (C) shows the spatial expression of all 523 detected DE genes grouped in three different clusters. From left to right: One cluster (1) contains genes which are lower expressed at the flg22 treatment area, two clusters contain genes with higher expression at the flg22 treatment spot in comparison to adjacent areas but with narrower (2) and broader (3) spatial expression. The yellow background in the plot indicates the flg22 treated area, the blue background indicates the water treated control area.

We compared both droplet spotting and syringe infiltration datasets (each dataset was collected 1 hour after flg22 exposure) for biological processes using GO-term enrichment. Both experiments produced a similar number of enriched GO-terms with 159 biological processes enriched in the droplet spotting dataset and 168 biological processes enriched in the infiltration dataset, with an overlap of 132 biological processes (83.0 % of the spotting dataset and 78.5 % of the infiltration dataset) between both datasets (see: **Additional_file2.xlsx**). The percentage of shared, enriched GO-terms indicated the presence of a similar plant response to flg22 in both experiments despite the difference in stimulus strength.

We measured the DE-genes with the 253 <u>fla</u>gellin rapidly <u>e</u>licited (FLARE) genes described by seedling and cell culture flg22 exposure experiment of Navarro et al. [18] (see: **Additional_file2.xlsx**). We found that the DE-genes of the droplet spotting experiment contained 24.90 % FLARE genes (63 of 253 genes). These consisted of 32 FLARE genes associated with signal transduction, 11 genes associated with roles in signal perception, 14 with known or putative roles as effector proteins and 9 FLARE genes identified by Navarro et al. as ‘other’ FLAREs (see: **Additional_file2.xlsx**). We found a slightly higher number in the infiltration experiment: 80 DE genes were shared with the 253 FLARE genes (31.62 %) with 39 genes associated with signal transduction, 16 genes in signal perception, 15 genes with known or putative roles as effector proteins and 11 genes with other functions (see: **Additional_file2.xlsx**). Interestingly the % of known FLARE and the number of enriched biological processes was higher in the infiltration than the spotting experiment (possibly suggesting other processes are triggered by infiltration).

We were interested in the spatial expression patterns of the 63 shared FLARE genes between our droplet spotting experiment and the Navarro et al. dataset. For this we visualised the expression patterns of the FLAREs across the studied leaf area. All 63 FLARE genes showed high expression levels at the area of flg22 exposure in comparison to adjacent leaf squares (Figure 5**B**). To study the expression profiles of the remaining 460 DE genes identified in the flg22 droplet spotting experiment, we affinity propagation clustered [21] these DE genes based on their spatial expression patterns and visualised the expression profile of each cluster (**Figure 4C**). We identified three gene clusters: two of the three clusters contained genes with higher expression levels at or adjacent to the area of flg22 treatment and one cluster contained a group of lower expressed genes at the flg22 treated area (**Figure 4C**).

**Figure 5.**
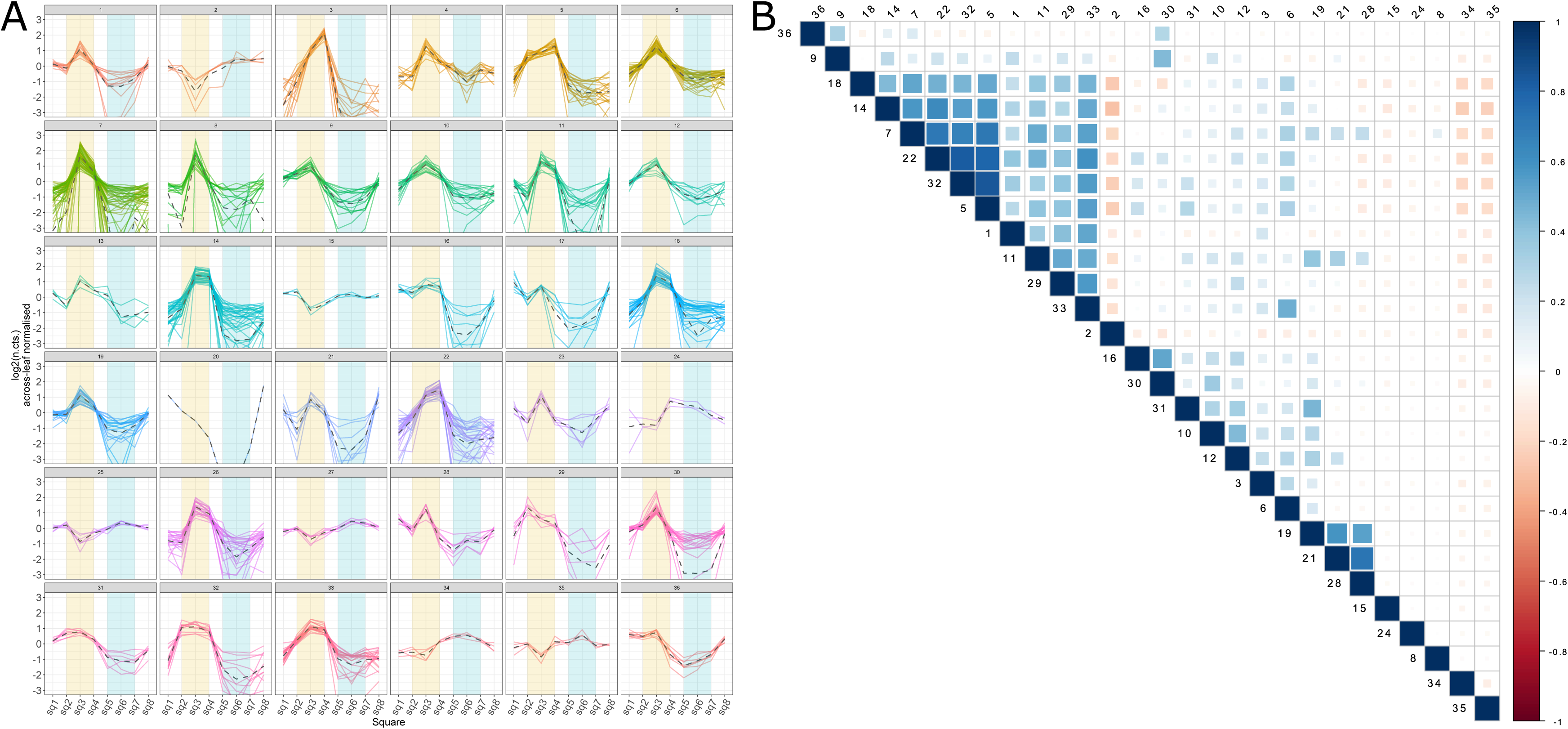
Unsupervised clustering of flg22 elicited DE genes and GO-term correlation matrix of the predicted clusters: Figure (A) shows the expression profiles of the 523 flg22 elicited DE genes grouped to 36 clusters precisely clustered according to their spatial expression pattern across the tested leaf area. Many of the clusters show differences in their induction profile at the site of flg22 deposition (yellow background) but also differences in expression at the water treated area (blue background) or the expression at the leaf boarders. Figure (B) shows the correlation analysis of the enriched GO-terms from the genes of the spatial clusters shown in (A) – 28 clusters grouped with hierarchical clustering for enriched GO-terms.

We analysed all clusters for enriched biological processes using GO-term analysis. We could not detect any enrichment in biological processes for the cluster containing the DE genes which were lower expressed at the flg22 site. The two clusters with expression peaks at the site of flg22 stimulation however enriched 135 and 122 biological processes. Of all biological processes 100 were shared between both clusters and 35 as well as 22 biological processes unique for each cluster respectively (see: **Additional_file1.docx** and **Additional_file2.xlsx**). This suggests that biological processes (host responses) could be associated with the spatial expression profiles of their corresponding genes. To study this further we used the affinity propagation clustering algorithm [21] to determine the number of clusters without a specified cluster number preference value. This grouped the 523 DE genes into 36 more tightly resolved spatial expression clusters, comprising 35 clusters with between three and 57 genes and a single cluster containing only one gene (**Figure 5A**). 28 of 35 clusters were enriched for biological processes (see: **Additional_file3.csv**). To test if different spatial expression patterns enrich different biological processes, we correlated all multi-gene clusters based on the presence / absence of all enriched GO-terms. We saw little overlap in biological processes between clusters, indicating that each spatial expression cluster enriched slightly different GO-terms. (**Figure 5B** and **Additional_file3.csv**).

### Characterisation of spatial regulatory elements

We characterised the expression patterns of the 36 obtained clusters in **Figure 5**. 11 clusters (1, 2, 6, 8, 12, 15, 23, 25, 28, 30, 35) showed a peak of higher expression at the site of flg22 perception. 14 clusters (3, 5, 7, 10, 11, 14, 18, 19, 21, 22, 26, 29, 32, 33) indicated spatially elevated gene expression patterns with higher expression also at sites adjacent to the area of flg22 perception. The remaining 11 clusters showed less clear expression profiles (**Additional_file2.xlsx**).

To identify plant regulatory elements that are potentially involved in PAMP perception and signal propagation to adjacent areas, we selected DE genes belonging to the flg22 locally and adjacently elevated clusters, and then filtered the genes for the TAIR-10 [22] GO-terms ‘receptor’ and ‘transcription’. This included the leucine-rich repeat receptor like kinase (LRR-RLK) *RLK7* (cluster 1) which was locally elevated, whereas the LRR-RLK *CERK1* and the serine/threonine-protein kinase *PSKR1* while strongly elevated at the area of flg22 perception were also broadly expressed throughout all sites (cluster 7). We detected a larger set of 48 DE genes associated with transcriptional processes. WRKY (15 genes), ERF (8 genes) and MYB (4 genes) transcription factor family members [23–25] were the most abundant in our dataset.

To start to understand the possible gene regulatory network controlling this spatial expression we used the TF2Network software [26] to search for putative regulatory interactions between these 48 DE genes i.e. by transcription factor binding. The resulting gene networks are built from genes with at least one target and a q-value < 0.01 (see **Additional_file1.docx**). These linked 4 transcription factors of which all belonged to the WRKY family (*WRKY11, WRKY15, WRKY17* and *WRKY47)* to 388 other DE genes indicating a possible regulatory network (TF2Network authors suggest their tools has a very low false positive rate, whilst being sensitive enough to detect 75-92% of correct links).

Of the detected transcription factors *WRKY17* (cluster 4) and *WRKY47* (cluster 8) were associated with local expression patterns, whereas *WRKY11* (cluster 10) and *WRKY15* (cluster 10) showed spatially wider expression.

## Discussion

The ability to profile gene expression patterns in small specific areas without bulk sequencing provides access to lower level transcripts, especially tissue and cell specific ones [3–5,28–30]. Spatial, low RNA-input transcriptomics methods allow deeper insights in how an organism develops and reacts to its environment than conventional “gross-scale” RNA-seq methods [4–6]. By combining rapid dissection with Grid-seq, we were able to reconstruct spatial transcriptional differences across organs and localised defence responses.

Although some specialised protocols are already available to profile transcriptomes from minute input amounts such as single-cells [2,10], or even nuclei [11,12], these detailed techniques do not retain the spatial information of a starting tissue and any time-consuming experimental procedures to preserve spatial data could induce experimental bias by altering the transcriptome. For spatial analyses in plants LCM [14,15] methods for fine-scale transcriptome analyses are available, however, these procedures are time-consuming and this limits the scale of the application. Large scale spatial analysis have been performed in the past [8], but still required bulk sampling of material by pooling multiple replicates. Methods relying on sample dissection and reaction-tube processing of tissue sections to sequencing libraries have already proven to be able to identify transcripts patterns in the zebrafish embryo [5] allowing to process multiple samples easily for modelling the transcriptome landscape of an entire organism. Recently Giacomello et al. [6] published a workflow to blot the transcriptome landscape from permeabilised plant tissues by vertical diffusion onto a slide containing an array of barcoded primers and on slide library construction which maintained the mRNA’s location via the barcode. This method, for the first time in plants, allowed access to spatial transcriptome data in thin tissue slices of plant organs with a great level of resolution. However, gathering and optimisation of permeabilization conditions of thin tissue sections can be challenging especially if the tissue due to the volume and shape of the sample, are not suitable to be processed on an array and the workflow is only commercially accessible.

Plants grow in a microbiologically rich environment with their own microbiomes and even symbionts [31], but they are also attacked by pathogens, pests, herbivores and other biotic stresses [32]. As plants can’t move away from attacks they defend themselves using molecular and cellular biology responses, however overstimulation of these processes leads to stunted development and lower fitness [17]. As plants must balance the need to defend themselves against constant plant-microbial interactions and attack [17] we hypothesised that local attacks might be integrated into a plant-wide defence response decision. Yet there are no appropriate assays to measure the molecular and cell biology changes at the required resolution. Here, we demonstrate a novel micro-scale method to pursue spatial transcriptomics experiments in plants in an easy manner based on easily accessible methods in sequencing library construction and bioinformatics tools.

We developed a robust micro-spatial expression methodology that enables the creation of transcriptome level maps from very small amounts of any eukaryotic tissue. The Grid-seq workflow evolved by transferring elements from existing single-cell RNA-seq methods [33,34] from animal systems to plants and refining these methods for stable, low-cost generation of sequencing libraries from small amounts of RNA starting material. This allows the design of experiments in which spatial information is required but only small pieces of tissue can be obtained. In the process of method development, we tested and included several features to efficiently generate double stranded cDNA (ds-cDNA) with reduced PCR amplification in both ds-cDNA synthesis and subsequent amplification after Nextera tagmentation. We introduced sample specific barcodes in the Nextera amplification step to allow pooling of 2304 of samples per sequencing run. This optimisation altogether allowed us to construct sequencing libraries by hand for just £ 6.00 per library (see **Additional_file2.xlsx**) in comparison to £ 65.56 for an Illumina TruSeq library (RS-122-2001, Illumina) or £ 62.60 for a SMARTer PCR cDNA Synthesis Kit library (634926, TaKaRa).

In our benchmarking experiments we compared the Grid-seq workflow with the widely used Illumina TruSeq sequencing protocol and show that the Grid-seq method compares well with this common commercial RNA-seq protocol (see: **Additional_file1.docx**). We also show that Grid-seq can detect transcript level differences across 1D leaf sections in distinct leaf elements such as leaf margins or vascular tissues and that spatial mapping of transcript levels to specific sections of leaves is possible, which allows drawing of transcriptional expression profiles across tissues. This easily used, low cost protocol makes feasible experiments that require spatial transcriptome analysis.

To apply the Grid-seq method for studying biotic actions we challenged *A. thaliana* leaves with the bacterial peptide flg22, a conserved 22 amino acid sequence of the bacteria flagellin protein, which to the plant indicates an encounter with potentially pathogenic bacteria [35]. Plants recognize such potential threats as the pathogenic cell surface molecules, so called Pathogen-Associated Molecular Patterns (PAMPs), perceived by the plant Pattern Recognition Receptors (PRRs) on the plant cell surface [36]. This event initiates an intracellular plant signalling cascade leading eventually to immunity or disease [17,37]. In our experiments we could detect the triggering of immune and defence response related biological processes and show that the results obtained by RNA-seq are independently reproducible using qRT-PCR (see: **Additional_file1.docx**). We were able to find overlap in our data with already described flg22 elicited (FLARE) genes from a gross-scale experiment using a strong stimulus [18]. In comparative analyses of our dataset with the spatial expression patterns of the described FLARE genes we were able to identify genes which share similar spatial expression and are potential novel FLARE genes. Cluster based analysis of spatial expression data revealed sets of genes with highly similar expression profiles enriched in distinct biological processes; including FLARE genes to which we add new and increased expression resolution. Characterisation of spatial cluster expression profiles highlighted plant regulatory elements with local or spatially elevated expression levels and so potential short distance signal propagators upon flg22 stimulus.

In a proof of principle experiment applying of the oomycte *Albugo laibachii NC14* to *A. thaliana* leaves we also recovered both host and pathogen transcript data, indicating that dual-host pathogen experiments (between eukaryotic organisms) are possible (see: **Additional_file1.docx** and **Additional_file2.xlsx**).

## Conclusions

We find that Grid-seq allows reproducible spatial-transcriptome experiments in plant tissues. Proof of concept experiments highlighted the usability of the method in characterising spatial differences within organs, and changes induced by biotic stimuli of bacterial PAMPs. The workflow also compares well to widely used RNA-seq protocols such as Illumina TruSeq sequencing but can be conducted at a fraction (∼ 1/10th) of the library preparation costs of Illumina TruSeq. Furthermore, as our ds-cDNA synthesis method is based on single-cell technologies [33], even smaller sections could be processed while maintaining efficient library and sequencing results.

## Methods

### Plant growth

For our experiments we used 4 – 6 week old *A. thaliana* Col-0 plants that were grown in a controlled environment room with an 8 hours light, 16 hour dark cycle at a constant temperature of 22 °C and 70 % humidity.

### Flg22 exposure experiments

Before flg22 treatment experiments, we transferred the plants from the controlled environment room to a laboratory working bench (room with constant light exposure and temperature). To elicit plant responses with flg22 we either syringe infiltrated [38] the peptide or spotted a droplet of flg22 on a leaf using a pipette.

To produce small, local infiltration spots we used a 1 ml syringe (BS01T, R&L Slaughter Ltd, Basildon, UK) loaded with 500 nM flg22 peptide solution. By application of mild pressure on the plunger of the syringe when infiltrating we produced an approximately 2 - 3 mm diameter infiltration spot on the left-hand side of a leaf. In parallel to flg22 infiltration we produced an infiltration series with DNase/RNase-free water as control. The plants were subsequently incubated on the laboratory working bench for 1 hour until sampling.

For the flg22 spotting experiment we loaded a 1 µl droplet of 500 nM flg22 on the abaxial surface of a leaf using a pipette (diameter approximately 1 mm). The flg22 was pipetted onto the left half of the leaf and a 1 µl droplet of the water control droplet spotted on the right half of the leaf. After spotting the plants were incubated for 1 hour on the laboratory bench before sampling.

### Leaf sectioning and sample harvesting

We used single margin razor blades (T586, Agar Scientific Ltd., Stansted, UK) to cut leaves into approximately 1 mm^2^ small leaf squares. To create a clean surface for cutting we used the pealed, non-sticky paper cover of a 96-well plate seal (AB0580, Thermo Fisher Scientific, Waltham, USA). With a previously in RNaseZAP (AM9780, Thermo Fisher Scientific, Waltham, USA) washed and air-dried forceps (T083, TAAB Laboratories Equipment Ltd, Berks, UK) we transferred each leaf square immediately after cutting into a well of a 96-well plate (E1403-0100-C, Starlab, Milton Keynes, UK) which we had pre-cooled on a 96-well metal block in dry ice (−70 °C), or alternatively, a dry-ice cooled 1.5 ml tube (10051232, Fisher Scientific, Loughborough, UK). The sample wells of 96-well plates were sealed using domed PCR cap strips (AB0602, Thermo Fisher Scientific, Waltham, USA). Post harvesting the samples were stored at −80 °C until use.

### Leaf sample lysis and preparation for mRNA extraction

To lyse the leaf samples stored in 1.5 ml tubes we first added 10 µl lysis buffer composed of 100 mM Tris-HCl pH 7.5 (BP1757, Fisher Scientific, Loughborough, UK), 500 mM LiCl (L7026, Sigma Aldrich, St. Louis, USA), 10 mM EDTA pH 8.0 (E7889, Sigma Aldrich, St. Louis, USA), 1% LiDS (L4632, Sigma Aldrich, St. Louis, USA), 5 mM DTT (18064014, Thermo Fisher Scientific, Waltham, USA) to each sample immediately after removing the sample tube from the cold storage.

We subsequently ground the leaf sections in lysis buffer using polypropylene pestles (Z359947, Sigma Aldrich, St. Louis, USA), which, before use, were washed with RNaseZAP (R2020, Sigma Aldrich, St. Louis, USA, three times with 80% ethanol (32221, Sigma Aldrich, St. Louis, USA) rinsed with UltraPure DNase/RNase-Free Distilled Water (10977049, Thermo Fisher Scientific, Waltham, USA) and air-dried after washing. After sample lysis we transferred the lysate to an ice-cooled 96-well plate and continued with the mRNA extraction.

Samples stored in 96-well plates were lysed by using 1 mm diameter grade 1000 hardened 1010 carbon steel ball bearings (Simply Bearings Ltd, Leigh, UK). For this, before use of the ball bearings, we treated a bulk batch sequentially with RNaseZAP and DNA AWAY, after this washed the ball bearings three times with 80% ethanol and transferred them to sterile screw-cap 2.0 ml tubes (E1420-2341, Starlab, Milton Keynes, UK) and heat dried with a slightly loosened lid on a 95 °C heating block (N2400-4001, Starlab, Milton Keynes, UK).

To lyse the collected leaf samples stored in a 96-well plate, we transferred the 96-well plate to a dry ice temperature cooled 96-well metal block. We carefully opened the domed PCR cap lids to avoid sample spillage and added the 4 – 6 (room temperature) ball bearings to each sample well. After this we transferred 10 µl lysis buffer to each well and re-sealed the plate with new domed PCR cap lids, and immediately proceeded to the 2010 Geno/Grinder (SPEX SamplePrep, Stanmore, UK) disrupting the samples for 30 seconds at 1750 rpm. We gathered the sampled using a centrifuge (Centrifuge 5910 R, Eppendorf UK Ltd, Stevenage, UK) for 10 seconds at 2000 x rcf. A strongly green-coloured solution without any remaining solid leaf material indicated good sample lysis. If satisfactory sample lysis was not achieved, we disrupted the samples again for another 10 seconds on the 2010 Geno/Grinder at 1750 rpm and centrifuged for 30 seconds at 2000 x rcf. We immediately transferred the lysis solutions into a new 96-well plate using a 10 µl multichannel pipette. After transfer of the lysis solutions, we stored the new 96-well plate on ice, discarded the 96-well plate containing the ball bearings and proceeded immediately with mRNA extraction. For a laboratory compatible version of the workflow see: **Additional_file1.docx**.

### Leaf mRNA purification

The leaf tissue mRNA was purified using 1 µl NEBNext Poly(A) mRNA Magnetic Isolation Module oligo-dT(25) beads (E7490, New England Biolabs Ltd, Hitchin, UK) per extraction. Previous to the extraction the required volume of oligo-dT(25) magnetic beads was washed twice in 200 µl lysis buffer on a DynaMag-2 Magnet rack (12321D, Thermo Fisher Scientific, Waltham, USA) and resuspended in 10 µl lysis buffer for each 1 µl oligo-dT(25) beads input volume. The beads were mixed by a quick vortex and 10 µl of the resuspended beads were transferred to each well of the 96-well plate containing the lysis solutions. The wells were sealed with domed PCR cap strips, the 96-well plate vortexed briefly and attached to a tube rotator (444-0502, VWR International Ltd, Luterworth, UK) with adhesive tape. After 10 minutes rotation we collected the lysis solution at the bottom of the wells by spinning the plate for 10 seconds at 2000 x rcf and pelleted the oligo-dT(25) magnetic beads on a 96-ring magnetic plate (A001219, Alpaqua, Beverly, USA). Using a multichannel pipette we washed the oligo-dT(25) magnetic beads twice with 50 µl Wash Buffer A (10mM Tris-HCl pH7.5, 0.15M LiCl, 1mM EDTA, 0.1% LiDS) and once with Wash Buffer B (10mM Tris-HCl pH 7.5, 0.15M LiCl, 1mM EDTA). After washing we centrifuged the plate for 10 seconds at 2000 x rcf to collect the remaining Wash Buffer B at the bottom of the tube, pelleted the oligo-dT(25) magnetic beads on a magnet and removed the remaining volume of Wash Buffer B with a multichannel pipette. The oligo-dT(25) beads were resuspended immediately in 8 µl DNase/RNase-Free water, incubated for 2 minutes at 80 °C on a G-Storm GS1 thermal cycler (G-Storm, Somerton, UK), then immediately pelleted on a 96-ring magnetic plate to elute the mRNA off and separate from the oligo-dT(25) beads. The solutions containing the purified mRNA were immediately transferred to a new 96-well plate, which was placed in a −80 °C freezer until needed. For a laboratory compatible version of the workflow see: **Additional_file1.docx**.

### Double-stranded cDNA synthesis reaction

For ds-cDNA synthesis we used a protocol based on the template switching mechanism of the reverse transcriptase enzymes [39]. Briefly: 2.50 µl extracted mRNA was mixed with 2 µl 5x First Strand buffer (18064014, Thermo Fisher Scientific, Waltham, USA), 1 µl 10 mM dNTPs (10297018, Thermo Fisher Scientific, Waltham, USA), 1 µl 5’-biotinylated 10 µM STRT-V3-T30-VN oligonucleotide: 5’-/5Biosg/TTAAGCAGTGGTATCAACGCAGAGTCGACTTTTTTTTTTTTTTTTTTTTTTTTTTTTTVN-3’ (Integrated DNA Technologies, Leuven, BE), 1 µl 20 mM DTT (18064014, Thermo Fisher Scientific, Waltham, USA), 0.10 µl 40 U/µl RNase Inhibitor (M0314S, New England Biolabs Ltd, Hitchin, UK), 0.25 µl 10 µM template switching oligo 5’-AAGCAGTGGTATCAACGCAGAGTGCAGUGCUTGATGATGGrGrGrG-3’ (Integrated DNA Technologies, Leuven, BE), 0.30 µl 200 U/µl SuperScript II Reverse Transcriptase (18064014, Thermo Fisher Scientific, Waltham, USA), 0.30 µl 100 µM MnCl2 (M1787, Sigma Aldrich, St. Louis, USA) and 1.55 µl DNase/RNase-Free water to a total reaction volume of 10 µl. The reverse transcription reaction was run in a G-Storm GS1 thermal cycler for 90 minutes at 42 °C with additional 10 minutes at 72 °C to inactivate the reverse transcriptase. After reverse transcription we immediately added 2 µl RNase H (M0297S, New England Biolabs Ltd, Hitchin, UK) diluted to 0.5 U/µl (5 U/µl stock concentration) to the reaction and incubated the reaction in the GS1 thermal cycler for 30 minutes at 37 °C. The RNase H treated reactions were purified using a 0.83x (10 µl) AMPure XP bead ratio (Beckman Coulter, High Wycombe, UK) and eluted in 18 µl 1x TE buffer. After this step we added 5 µl 5x Kapa HiFi PCR buffer (KK2102, KAPA BioSystems, Wilmington, USA), 0.75 µl 10 mM dNTPs, 0.75 µl 10 µM PCR+G primer 5’-GAAGCAGTGGTATCAACGCAGAGT-3’ (Integrated DNA Technologies, Leuven, BE) and 0.50 µl 1 U/µl Kapa HiFi polymerase (KK2102, KAPA BioSystems, Wilmington, USA) to the cleaned ds-cDNA resulting in a total reaction volume of 25 µl and amplified the ds-cDNA in a G-Storm GS1 thermal cycler according to the following programme: (1) 3 minutes at 94 °C, (2) 17 cycles with 30 seconds at 94 °C, 30 seconds at 63 °C and 1 minute 30 seconds at 72 °C, (3) a final elongation step for 5 minutes at 72 °C. The amplified libraries were purified using a 1x (25 µl) AMPure XP bead ratio and eluted in 20 µl 1x TE buffer. The ds-cDNA libraries could be stored at this point in a −20 °C freezer. Before continuing with Illumina sequencing library preparation, we measured the ds-cDNA library concentrations with the Qubit 2.0 Fluorometer (Thermo Fisher Scientific, Waltham, USA) dsDNA HS Assay Kit reagents (Q32854, Thermo Fisher Scientific, Waltham, USA) and also assessed the size distributions of randomly picked libraries on an Agilent Bioanalyser (G2939BA, Agilent Technologies, Stockport, UK) using the Agilent High Sensitivity DNA Kit (5067-4626, Agilent Technologies, Stockport, UK).

At later stages we modified the ds-cDNA synthesis integrating elements of the Smart-seq2 protocol [40]. The reverse transcription reactions of Grid-seq-1.0 (as described above) and Smart-seq2 were already highly similar, but Smart-seq2 had proven to require less hands-on time than the Grid-seq-1.0 reverse transcription workflow. Grid-seq-1.1 uses the Smart-seq2 ds-cDNA synthesis with minor modifications, briefly: 2.50 µl extracted mRNA were combined with 1 µl 10 µM Smart-seq2 Oligo-dT30VN (5’-AAGCAGTGGTATCAACGCAGAGTACTTTTTTTTTTTTTTTTTTTTTTTTTTTTTTVN-3’, Integrated DNA Technologies, Leuven, BE) and 1 µl 10 mM dNTPS to a total volume of 4.5 µl. To anneal the Smart-seq2 Oligo-dT30VN we incubated the library for 30 seconds at 72 °C and snap-cooled the mixture on ice. The reverse transcription was conducted by adding the following reagents to the reaction with a final reaction volume of 10 µl: 2 µl 5x First Strand buffer, 2 µl 5 M betaine (B0300, Sigma Aldrich, St. Louis, USA), 1 µl 1 M MgCl2 (AM9530G, Thermo Fisher Scientific, Waltham, USA), 0.5 µl 100 mM DTT, 0.25 µl 40 U/µl RNase Inhibitor (2313A, Takara Clontech, Mountain View, USA), 0.10 µl 10 µM Smart-seq2 template switching oligo (5′-AAGCAGTGGTATCAACGCAGAGTACATrGrG+G-3′, Exiqon, Vedbaek, DK), 0.50 µl 200 U/µl SuperScript II Reverse Transcriptase, and 0.09 µl DNase/RNase-free water. We performed the reverse transcription reaction for (1) 90 minutes at 42 °C, (2) 15 cycles with 2 minutes at 50 °C and 2 minutes at 42 °C and finally (3) 15 minutes at 70 °C. After reverse transcription we added 12.50 µl 2x Kapa HiFi HotStart ReadyMix (KK2601, KAPA BioSystems, Wilmington, USA), 0.25 µl 10 µM Smart-seq2 IS-PCR primers (Integrated DNA Technologies, Leuven, BE) and 2.25 µl DNase/RNase-Free water to the reaction resulting in a total volume of 15 µl per reaction. Amplification was performed in a G-Storm GS1 cycler according to the programme: (1) 3 minutes at 98 °C, (2) 15 cycles with 20 seconds at 98 °C, 15 seconds at 67 °C and 6 minutes at 72 °C and a (3) final elongation step for 5 minutes at 72 °C. The PCR reactions were purified using a 0.65 x (9.75 µl) AMPure XP cleanup and eluted in 20 µl 1 x TE buffer. After cleanup we measured the ds-cDNA library concentrations with the Qubit 2.0 Fluorometer dsDNA HS Assay Kit reagents and loaded randomly selected libraries on the Agilent Bioanalyser using the Agilent High Sensitivity DNA Kit before continuing with Illumina sequencing library prepration. For a laboratory compatible version of the workflow see: **Additional_file1.docx**.

### Illumina library preparation from ds-cDNA

We prepared Illumina sequencing libraries using an Illumina Nextera (FC-121-1030, Illumina Cambridge, UK) based protocol with minor modifications: we exclusively used the Tagment DNA Enzyme 1 and the Tagment DNA Buffer and amplified the tagmented DNA with the Kapa 2G Robust Polymerase (KK5024, Sigma Aldrich, St. Louis, USA). We used custom Nextera barcodes that allow us to multiplex hundreds of samples (see: **Additional_file2.xlsx**) [41].

We reduced the costs of the library preparation by reducing the total tagmentation reaction volume to 5 µl (from 50 µl as recommended) with 1 ng ds-cDNA library input and using less enzyme. We performed a titration experiment of Tagment DNA Enyzme vs. 1 ng of selected ds-cDNA libraries aiming for Illumina sequencing libraries with a modal insert size distribution in the range of 400-500 bp with little short insert fragments and found that 0.1 µl Nextera enzyme was optimal.

The Nextera reactions were performed by combining 1 ng of ds-cDNA (air-dried over-night at room temperature in a drawer with the 96-well plate loosely covered to allow evaporation of liquid) with 2.5 µl 2 × Nextera buffer, 2.4 µl water and 0.1 µl Nextera enzyme on ice. The tagmentation plate was immediately transferred for 5 minutes at 55 °C on a G-Storm GS1 thermal cycler. Meanwhile we prepared a fresh 96-well plate with 2.0 µl 2.5 µM P5 and 2.0 µl 2.5 µM P7 custom multiplexing primers (see: **Additional_file2.xlsx**). After tagmentation we transferred the tagmentation reactions to the previously prepared 96-well plate containing the sequencing adapters (see above) and added the following to each well: 5.00 µl 5 x Kapa 2G Robust Buffer, 0.50 µl 10 mM dNTPs, 0.10 µl 5 U / µl Kapa 2G Robust Polymerase, 10.4 µl water to a total final volume of 25 µl.

Amplification was performed on a GStorm GS-1 cycler using the following program: (1) 3 minutes at 72 °C, 1 minute at 95 °C (2) 11 cycles of 10 seconds at 95 °C, 30 seconds at 65 °C, 2 minutes 30 seconds at 72 °C (2) a final elongation step for 2 minutes 30 seconds at 72°C. After amplification we purified the libraries using a 0.64x ratio (16 µl) AMPure XP beads, measured the library yields with the Qubit 2.0 Fluorometer dsDNA HS Assay Kit reagents and assessed the size distributions of randomly selected libraries on the Agilent Bioanalyser with the Agilent High Sensitivity DNA Kit. For a laboratory compatible version of the workflow see: **Additional_file1.docx**.

### Sample pooling and sequencing

For sequencing all library concentrations were determined using Qubit 2.0 Fluorometer using the dsDHA HS Assay kit reagents and pooled at equal molarity. The profile and concentration of the final library pool was assessed on the Agilent Bioanalyser using Agilent High Sensitivity DNA Kit Sequencing reagents. After this the pooled samples were shipped to the Earlham Institute for sequencing. Quality control and data demultiplexing was performed by the Earlham Institute Genomics Pipelines facilities. Samples were sequenced using Illumina HiSeq2500 50 base single-end rapid run sequencing for the *A. thaliana* wounding and *A. thaliana* flg22 infiltration datasets Illumina NextSeq500 75 base single-end for the *A. thaliana* untreated leaf dataset and Illumina HiSeq4000 150 base paired-end sequencing for the *A. thaliana* flg22 droplet spotting experiment.

### Data quality control and mapping

The sequencing reads were quality controlled using FastQC-0.11.5 [42]. After quality control we used cutadapt-1.17 [43] to trim low-quality bases (-q 20) and remove Oligo-dT, template switching oligos, primer and Illumina Nextera library preparation sequences (-n 5 -e 0.05 –overlap 10). We also removed sequences with less than 40 bases (--minimum-length 40) and sequences containing N’s (--max-n 0) from the dataset with cutadapt-1.17. After adapter and quality trimming we re-assessed the reads a second time with FastQC-0.11.5. We mapped the reads to the *A. thaliana* TAIR10 release 37 genome assembly using STAR-2.5.1b [44] default settings and assessed mapping scores, duplication levels, GC-bias and gene-body coverage after mapping with RSeQC-2.6.4 [45]. Reads were counted with HTSeq-count-0.6.0 [46] default settings.

### Differential-expression analysis, GO-term enrichment and plant metabolic pathway enrichment

Differential expression analysis was performed using DESEq2-1.20.0 [47] in the statistical language R-3.5.1 using the workflow described by Love et al. [48] but by pre-filtering the dataset for rows with less than 10 rather than 1 raw read counts. DE-genes were called with a q-value threshold < 0.05%.

Across leaf DE-gene expression plots were prepared using R-3.5.1; in brief: We imported all samples with DESeq2-1.20.0 and calculated a table with normalised expression values as in the workflow described by Love et al. [48]. Next, we calculated the average expression value of each gene in each leaf square across all biological replicates. As a final step we normalised the expression values of the leaf squares. For this we divided the mean expression value of each leaf square of a gene with the mean expression value across all leaf squares of the same gene. The log2 transformed plots of the so normalised data were generated using ggplot2-3.1.0 [49]. Affinity propagation clustering of the normalised expression tables was performed using the R-3.5.1 library apcluster-1.4.7 and a Pearson distance matrix of the normalised data for apclustK and apclust [21] default settings. The number of clusters was either empirically determined by continuously increasing the preferred cluster number in the apclustK function and visualising the expression profiles of the clusters using ggplot2-3.1.0 or determined without providing a cluster number preference value using the apclust function.

GO-term enrichment analysis on DE-genes was performed using the R-3.5.1 Bioconductor library ClusterProfiler-3.8.1 [50] with the settings (Statistical test: Hypergeometric test, Multiple testing correction: Benjamini & Hochberg False Discovery Rate correction, False Discovery Rate cutoff: 0.01) and the Bioconductor library org.At.tair.db-3.6.0 as organism database [51].

## Additional files

### Additional_file1.docx

Comparison of Grid-seq with Illumina TruSeq, qRT-PCR validation of DE gene expression, wounding time-series GO enrichment, *Albugo laibachii* spatial transcriptomics experiment, all wet-lab protocols of the Grid-seq workflow.

### Additional_file2.xlsx

Read numbers, ENA accessions, DE genes, GO accessions, clustering information, FLARE associations of experiments.

### Additional_file3.xlsx

GO-term enrichments of spatial affinity propagation clustering.

## Ethics approval and consent to participate

Not applicable.

## Consent for publication

This study abides by UK guidelines and legislation for plant science research.

## Availability of data and material

Sequencing reads for the Grid-seq vs. TruSeq comparison are available under the ENA study accession number PRJEB31337. Sequencing reads for the wounding experiment are available under the ENA study accession number PRJEB31311. Sequencing reads for the untreated leaf spatial transcriptomics experiment are available under the ENA study accession number PRJEB31314. Sequencing reads for the flg22 / water infiltration experiment are available under the ENA study accession number PRJEB31312. Sequencing reads for the flg22 / water droplet spotting experiment are available under the ENA study accession number PRJEB31313. Sequencing reads for the *Albugo laibachii* x *Arabidopsis*.*thaliana* spatial transcriptomics experiment are available under the ENA study accession number PRJEB31803.

## Authors’ contributions

M.G., W.V. and D.H. established protocols. M.G., W.V. and A.L. prepared sequencing libraries. M.G. analysed the data. I.M. advised on protocols and data analysis. M.G. and M.D.C. wrote the manuscript. W.V., M.G. and M.D.C. designed the study.

## Acknowledgements

We thank David Prince for providing the *A. laibachii* NC14.

