## Additional_file1 for "Spatially resolved transcriptomics reveals plant host responses to pathogens"

### 1.1 Comparison of RNA sequencing library methods

To examine experimental reproducibility between the Grid-seq workflow and a standard macro protocol, we experimentally compared Grid-seq with the Illumina TruSeq RNA Sample Prep v2 kit (henceforth referred to as Illumina TruSeq).

The cDNA synthesis step of Grid-seq is based on the Smart-seq2 [19] protocol and generates double stranded cDNA (ds-cDNA) rich in full length transcripts. Smart-seq2 was developed for single-cell NA-seq, with each cell containing minute (< 100 pg) amounts of target mRNA. In contrast the Illumina TruSeq kit requires much larger input amounts i.e. total RNA of ~1000 ng per reaction and is used to assay transcriptomes at a macro-scale. We were interested in comparability of results between the two methods and the advantages of higher resolution transcriptomics. For this experiment we detached a single leaf from an *A. thaliana* plant and submerged the leaf for 1 hour in a 500 nM flg22 or water solution (3 biological replicates). After 1 hour we extracted the mRNA from the samples and used the same isolated mRNA as input for either Illumina TruSeq (1000 ng total RNA each) or Grid-seq (1 ng total RNA each) libraries. To standardise the amount of data used for method comparison analysis we normalised the number of reads for each biological replicate using 1000-times random sub-sampling of the raw count files from 5.46 ± 1.29 million (M) mapped reads for the TruSeq libraries and 4.00 ± 0.37 M mapped reads for the Grid-seq libraries to 1.5 M mapped reads (script used for subsampling: <https://github.com/mgiolai/Grid-seq/blob/master/Grid-seq_random-HTSeq.py>). We separately analysed each subsampling round. We found similar numbers of genes for both methods: 16247 ± 39 for Grid-seq and 16402 ± 55 for Illumina TruSeq. We called the DE-genes of each sub-sampling round by comparing the flg22 exposed leaves with the water controls. We found 185 ± 31 higher, 65 ± 13 lower expressed DE genes for Grid-seq, and 130 ± 17 higher, 344 ± 45 DE-genes lower expressed for Illumina TruSeq.

To assess the reproducibility of both library preparation methods in a more realistic scenario, we also determined the number of detected genes and the number of DE genes without random subsampling. Among the Grid-seq replicates with 4.00 ± 0.37 M mapped reads we detected 17647 genes and among the Illumina TruSeq replicates with a higher number of 5.46 ± 1.29 M mapped reads we detected 18829 genes. We found 788 higher expressed and 348 lower expressed DE genes for Grid-seq and 1042 higher and 323 lower expressed DE genes for Illumina TruSeq. 1636 of all DE genes overlapped with both datasets (69.4 % overlap with Grid-seq DE-genes and 83.4 % overlap with Illumina TruSeq DE-genes). We tested the shared DE genes for enriched biological processes using GO-term analysis and detected 247 enriched biological processes of which many are associated with roles in the plant immune response (**Figure 1**).

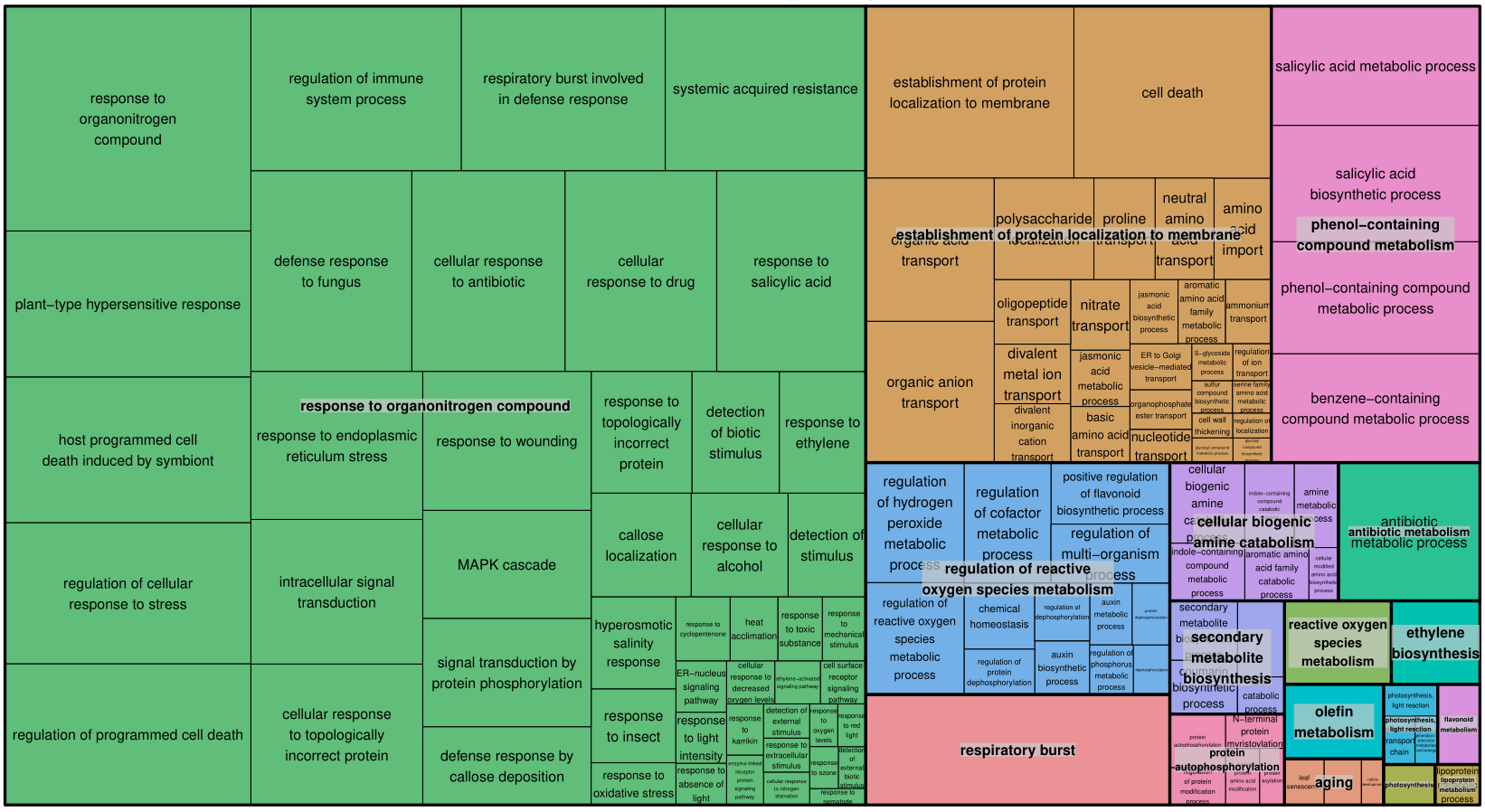

**Figure 1** Enriched GO-terms for the shared DE genes between the Illumina TruSeq and Grid-seq flg22 leaf treatment experiment visualised with REVIGO [2]. The size of each rectangle represents the absolute log10(q-value).

To observe how well library preparation methods and treatments correlate, we used a correlation analysis (pearson) on the normalised gene tables of both library preparation methods. The results of the correlation analysis are listed in **Table 1**. The correlation coefficients between Grid-seq and Illumina TruSeq datasets indicated that the library preparation methods compared well.

**Table 1** Pearson correlation analysis of Grid-seq (GS) and Illumina Truseq (TS) datasets.

| Comparison | Pearson Correlation coefficient (Min / Max) |
| --- | --- |
| flg22 (GS) | 0.94 / 0.95 |
| H2O (GS) | 0.96 / 0.97 |
| flg22 (TS) | 0.92 / 0.96 |
| H2O (TS) | 0.96 / 0.99 |
| flg22 (GS) vs flg22 (TS) | 0.92 / 0.96 |
| H2O (GS) vs H2O (TS) | 0.96 / 0.99 |
| TS ALL vs GS ALL | 0.92 / 0.99 |

### 1.2 Wounding time-series experiment

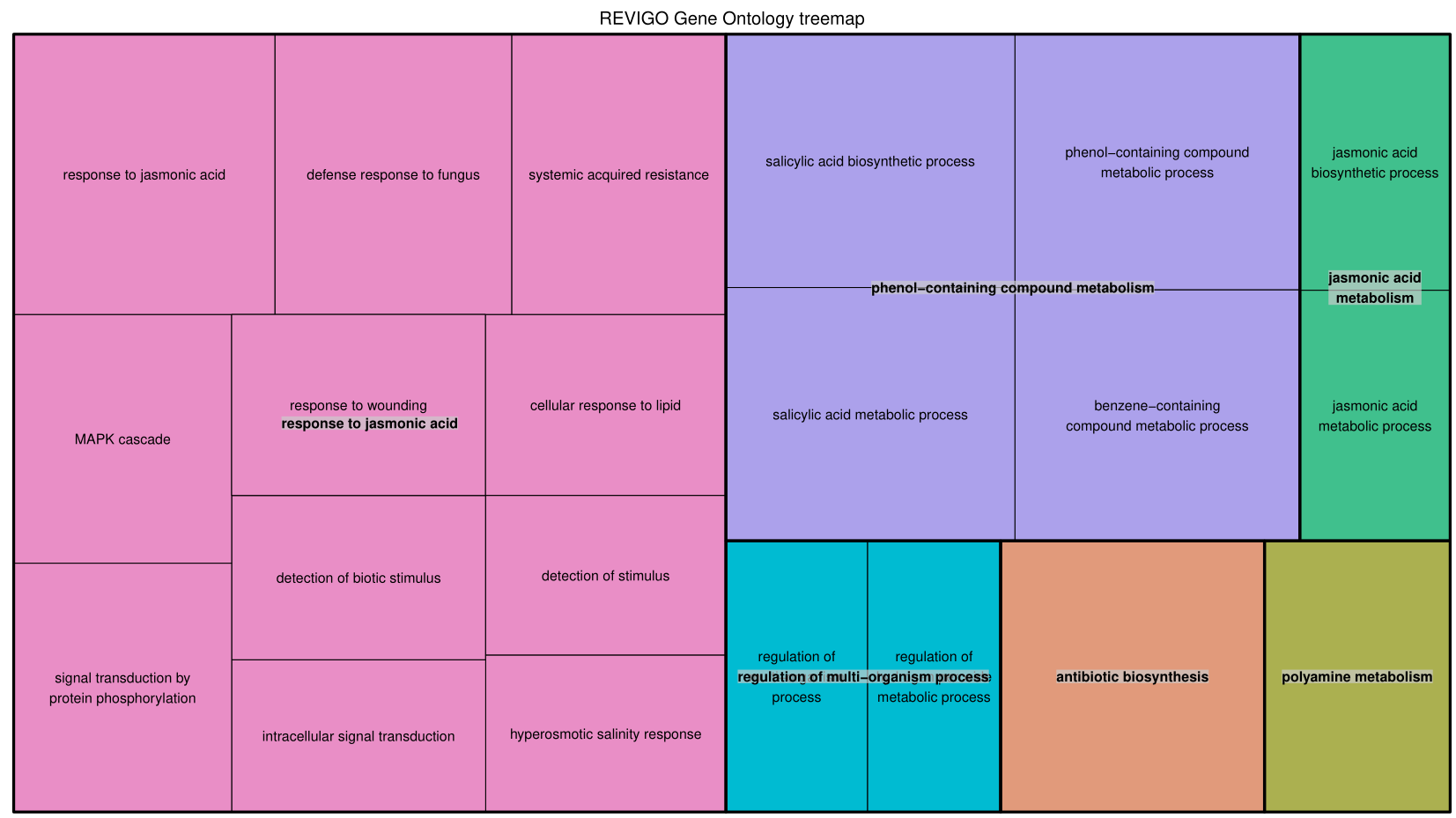

**Figure 2** Enriched GO-terms for the combined genes of the wounding time-series experiment visualised with REVIGO [2]. The size of each rectangle represents the absolute log10(q-value).

### 1.3 Validation of flg22 induced, local plant response genes using qRT-PCR

We tested the expression of DE-genes between the flg22 and water infiltration RNA-seq datasets using qRT-PCR. We selected 6 DE-genes upregulated at the flg22 infiltration spot in comparison to the water control over a range of log2-fold change values from 3.8 to 1.3 and q-values from 0.002 to 0.05 and repeated the infiltration experiment by comparing 8 localised flg22 infiltration spots with 7 water infiltration spots. The 6 selected DE-genes: transcription factor *WRKY30* (AT5G24110, log2-fold change=3.8, q-value=0.004), NAD(P)-binding protein gene *SDR5* (AT2G47140, log2-fold change=2.2, q-value=0.002), pathogenesis-related PR-6 proteinase inhibitor family member (AT2G38870, log2-fold change=2.1, q-value =0.043), putative LRR-RLK family member *IOS1* (AT1G51800, log2-fold change=1.4, q-value=0.033), a member of the UDP-Glycosyltransferase superfamily (AT1G05675, log2-fold change=1.6, q-value=0.005) and a hypothetical protein gene (AT2G25735, log2-fold change=1.3, q-value=0.05). We calculated the 2^-dCt^ values of the flg22 and water datasets by using the two house-keeping genes *PEX4* and *PTB1* for normalisation. All 6 selected DE-genes were verified -by statistically significant higher expression detected by qPCR in the flg22 exposed samples comparison to the water controls (**Figure 2**).

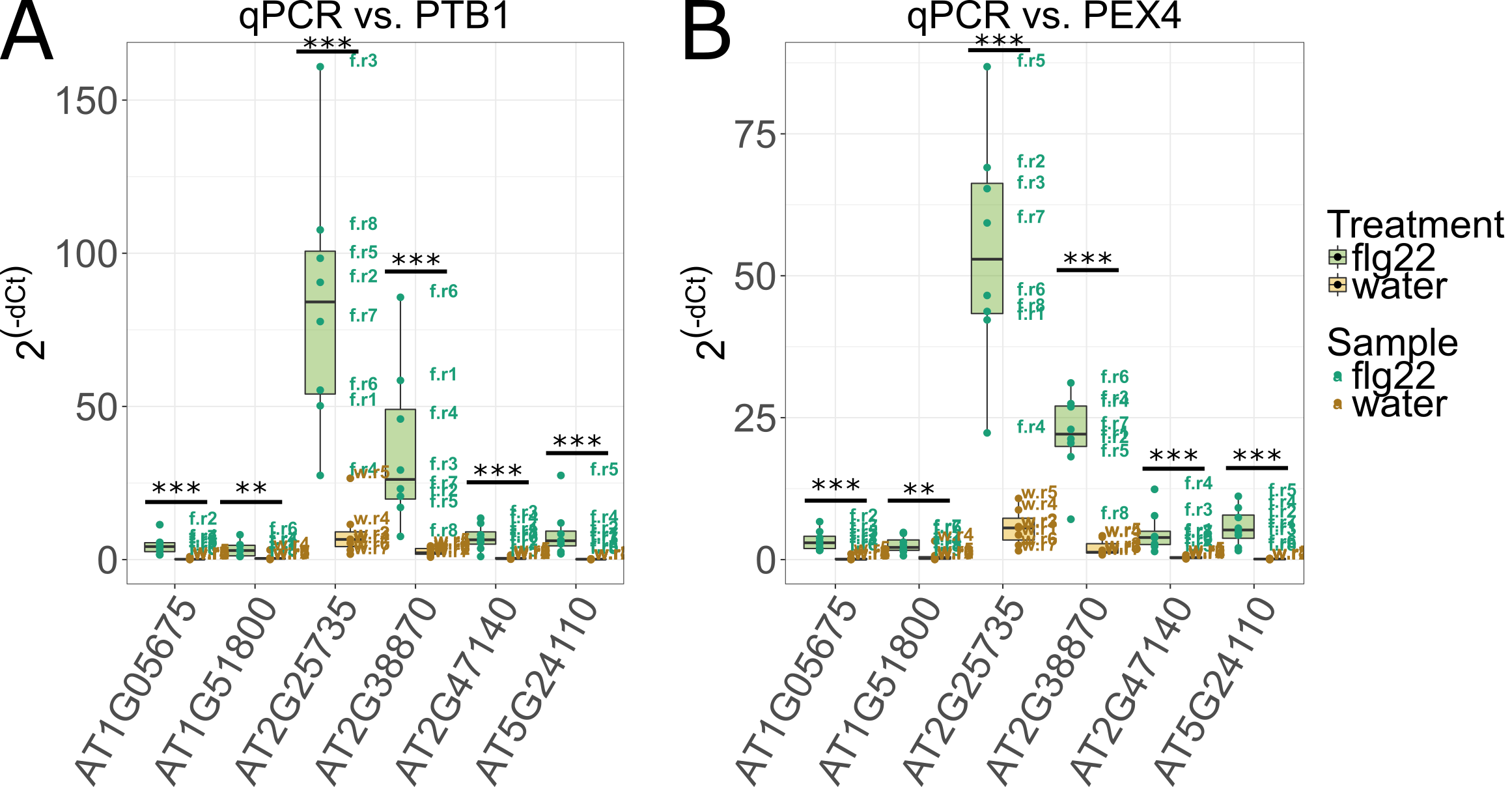

**Figure 3 qRT-PCR validation of flg22 responsive DE-gene expression:** To verify differential gene expression upon flg22 treatment we selected six genes. We assessed the expression levels of the selected genes in 8 biological replicates of flg22 infiltration and 7 biological replicates of the water infiltration controls. Image (A) shows the 2^-dCt^ of flg22 infiltrated samples (green) and water infiltrated samples (yellow) normalised to the housekeeping gene PTB1. Image (B) was produced by normalising the data to the housekeeping gene PEX4. All genes which are called as DE in the Grid-seq method are also statistically significant higher expressed in the qRT-PCR experiment (* = p ≤ 0.05, ** = p ≤ 0.01, *** = p ≤ 0.005; t-test).

#### 1.3.1 qRT-PCR reaction setup

For qRT-PCR reactions we quantified the ds-cDNA libraries with the Qubit 2.0 Fluorometer dsDNA HS Assay Kit reagents. Based on this quantification we diluted each ds-cDNA library to 100 pg/µl in 1x TE buffer and transferred 1 µl (equals 100 pg ds-cDNA) diluted ds-cDNA library to a 384-well qPCR plate (4TI-0381, 4titude Ltd, Wotton, UK) with an electronic Eppendorf Xstream pipetter (Eppendorf, Wesseling, Germany). After this we added 1.2 µl of a polymerase / primer master-mix composed of 1 µl 2x KAPA SYPR FAST Universal master mix (KK4601, Sigma Aldrich, St. Louis, USA), 0.04 µl 10 µM gene specific forward primer (Integrated DNA Technologies, Leuven, BE), 0.04 µl 10 µM gene specific reverse primer (Integrated DNA Technologies, Leuven, BE) and 0.12 µl DNase/RNase-free water to each reaction, also with the Eppendorf Xstream pipetter. The plates were sealed with qPCR plate seals (4ti-0560, 4titude Ltd, Wotton, UK), centrifuged for 4 minutes at 2000 x rcf (Centrifuge 5910 R, Eppendorf UK Ltd, Stevenage, UK) and transferred to the LightCycler 480 Instrument (Roche, Burgess Hill, UK) for qPCR cycling according to the following programme: (1) 3 minutes at 95 ⁰C, (2) 45 cycles with 3 seconds at 95 ⁰C (4.8 ⁰C/s ramp rate), 20 seconds at 60 ⁰C (2.5 ⁰C/s ramp rate) and single acquisition mode for each cycle, (3) melting curve assessment with 5 seconds at 95 ⁰C (4.8 ⁰C/s ramp rate), 1 minute at 65 ⁰C (2.5 ⁰C/s ramp rate) followed by ramping to 97 ⁰C (0.11 ⁰C/s ramp rate) in continuous acquisition mode with 5 acquisitions per ⁰C. Transcript Cp values were determined using the LightCycler 480 Software release 1.5.0 ‘Abs Quant/2^nd^ Derivative Max for All Samples’ analysis method, melting curves were calculated using the ‘Tm Calling for All Samples’ analysis method. Further analyses were performed with Microsoft Excel (the qRT-PCR oligonucleotides are listed in the **Table 2**).

**Table 2 Olignucleotides used for qRT-PCR.**

| Name | Sequence (5‘-3‘) |
| --- | --- |
| AT1G05675-F | ACAAGCCCTCTCCGCCGTAC |
| AT1G05675-R | GGTCTTCTGATCGTTCCTGGCCT |
| AT1G51800-F | ACCGCAAATAGTTCACCGCGAC |
| AT1G51800-R | TGGAAAAGACCGTGAAAGCCCGA |
| AT2G25735-F | GCAACACCAACGAAGAGAAAGGC |
| AT2G25735-R | AGTGGAAGAAGCACGAGAAGGCA |
| AT2G38870-F | GGTGACTATGCGGCCGTGGT |
| AT2G38870-R | GCACCTGAAGTCTGCGGTCAC |
| AT2G47140-F | CGATCGCCATTAACCTCCGCG |
| AT2G47140-R | GTGCAAACGATGGAGCCGCG |
| AT5G24110_F | ACTACTCCGGCGAACTTGAC |
| AT5G24110_R | GCATTGACTTCTTCGAACTCTTG |
| AT5G25760_F | TGCAACCTCCTCAAGTTCG |
| AT5G25760_R | CACAGACTGAAGCGTCCAAG |
| AT3G01150_F | ACCCATCATCTGATCCCAAC |
| AT3G01150_R | AACATGGAAATACTCCCCATTG |

### 1.4 Early plant response genes of local, fine-scale flg22 stimulation (flg22 droplet spotting)

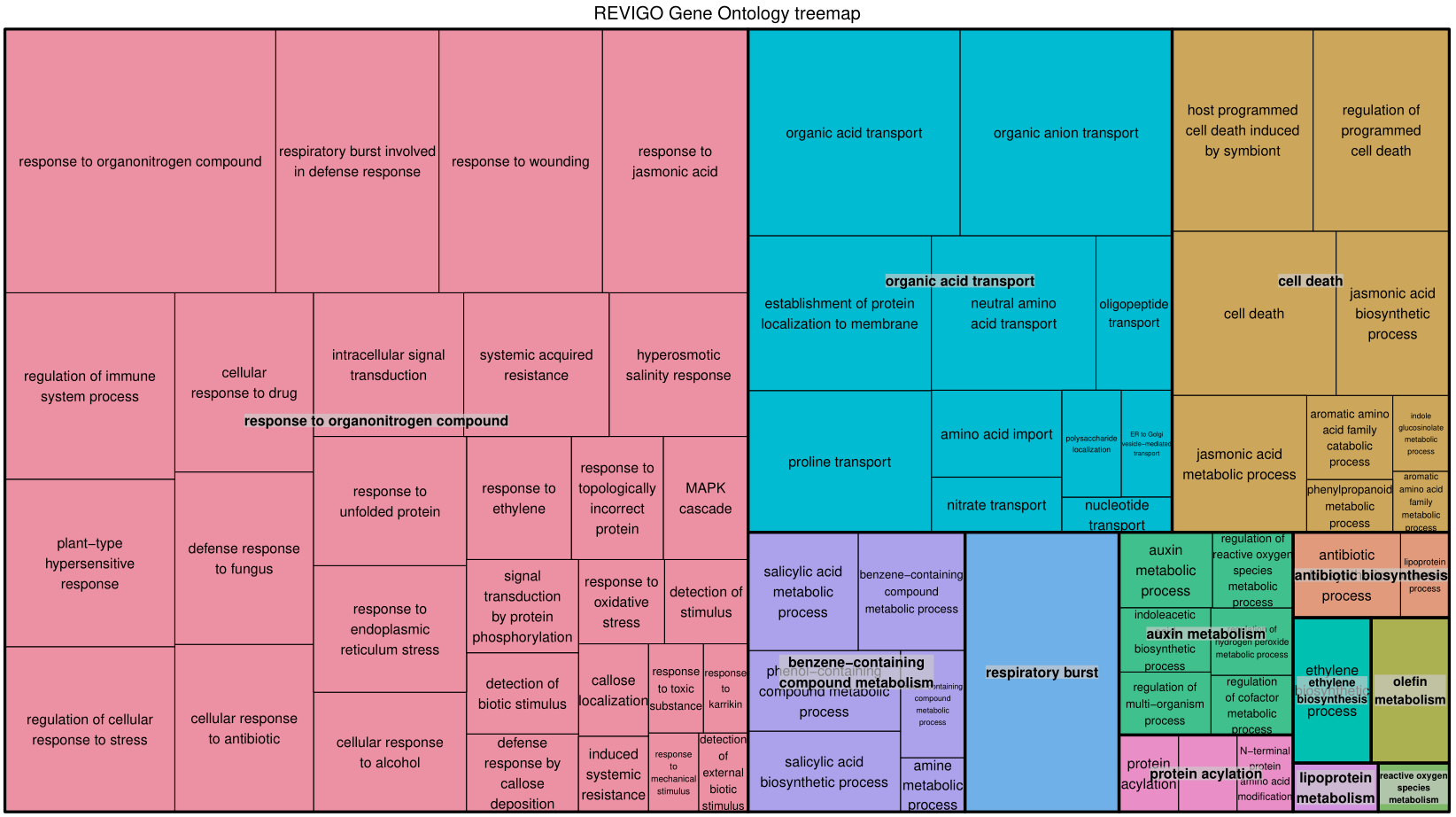

**Figure 4** REVIGO [2] plot of GO-terms enriched using the DE-genes of spatial cluster 2 (of 3 total clusters). The size of each rectangle represents the absolute log10(q-value).

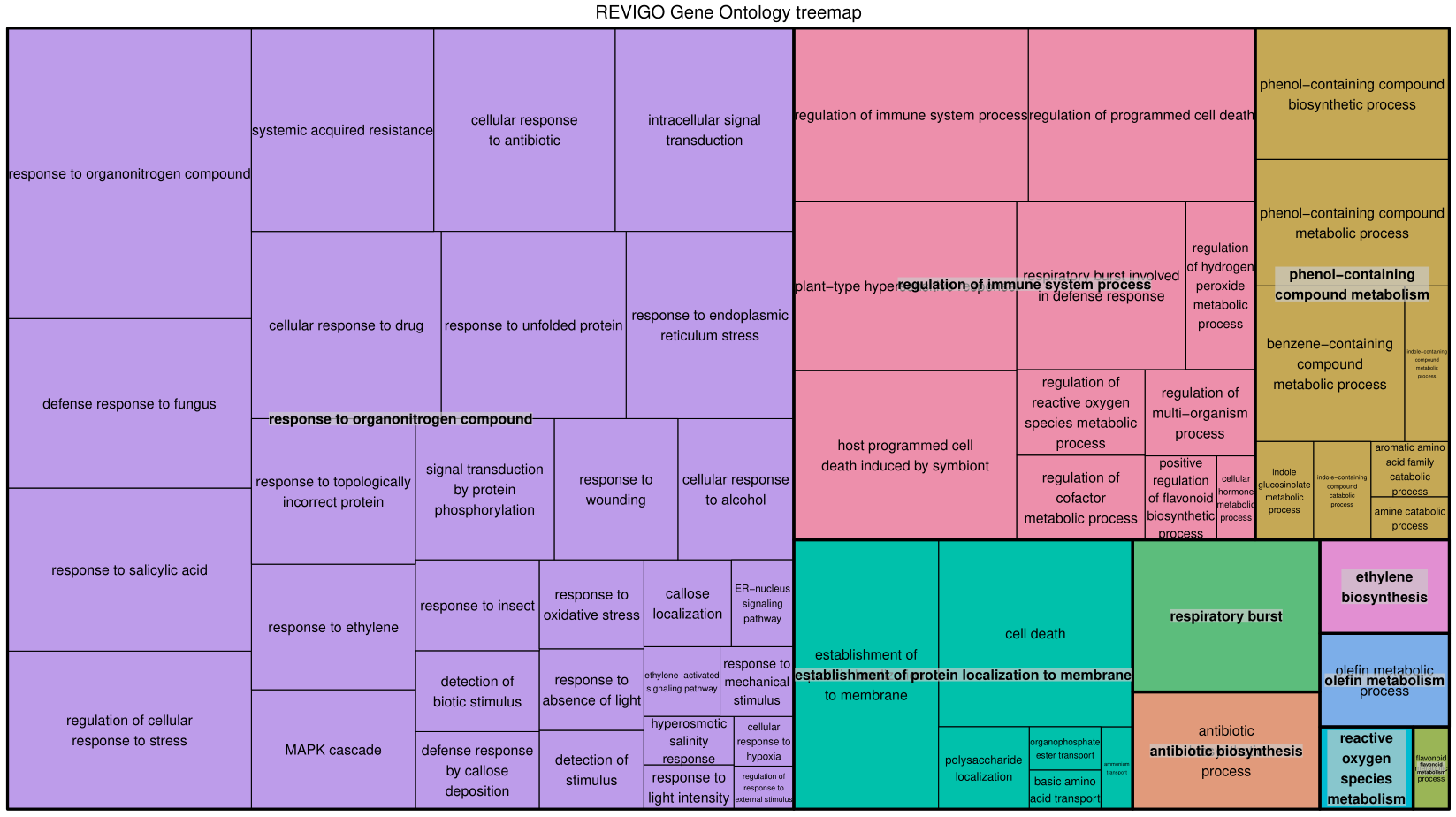

**Figure 5** REVIGO [2] plot of GO-terms enriched using the DE-genes of spatial cluster 3 (of 3 total clusters). The size of each rectangle represents the absolute log10(q-value).

### 1.5 *Albugo laibachii* NC14 dual host-pathogen RNA-seq experiment

We exposed a ~ 1mm^2^ large abaxial leaf-area of 4-week-old *A. thaliana* Col-0 plants in two biological replicates to *A. laibachii* NC14 sporangia (5.6 E6 sporangia / 1 ml). With control plants (two biological replicates) we spotted water on the same leaf area. The plants were incubated for 5 days in in a growth cabinet with an 8 hour light (180 µmol/m^2^ s) at 20 ⁰C and a 16 hour dark at 14 ⁰C cycle. We extracted the oomycete exposed area after 5 days as square-3 and laterally moving to the left, square-1 and square-2 and the right square-4 and square-5 as non-vascular tissue areas. The libraries were prepared with the Grid-seq-1.0 workflow and sequenced for 50 single-end reads on an Illumina HiSeq2500 instrument. The reads were mapped to both genomes (*A. thaliana* TAIR10 release 37 genome and *Albugo laibachii* ENA1) as described in the manuscript. For filtering reads into SAM files which mapped (a) better (determined by the alignment score in the SAM file) to one of the two genomes or (b) uniquely to a genome we used the ‘Grid-seq_compare-SAM.py’ script (https://github.com/mgiolai/Grid-seq/blob/master/Grid-seq_compare-SAM.py). We then counted the reads in the filtered SAM files and used this output for all further analyses.

We analysed the numbers of mapping reads in the controls and oomycete exposed sections and counted 8161, 4823, 312021 and 5417 reads mapping to *A. laibachii* in the square-3 tissue section of control 1 (ctrl1) and – 2 (ctrl2), as well as oomycete exposed replicate 1 (oom1) and – 2 (oom2) respectively. As the number of reads in oom2 was similar to the number of reads in the controls, we assumed that the infection of this replicate might not have been as efficient as in the oom1 sample. Therefore, to control for the presence of *A. laibachii* transcripts across the 1 D leaf map, we extracted the genes of oom1 with higher, normalised DESeq2 expression scores (≥ 10) and prepared an expression plot of the transcripts across the leaf squares. This resulted in filtering of 663 genes and showed a clear enrichment for *Albugo laibachii* presence in square-3 (**Figure 6**). When calling DE genes between the square-3 areas of the oomycete exposed leaves and the water controls we detected 1054 higher and 763 lower expressed genes. Enriched GO-terms (biological processes) can be observed in **Figure 7**.

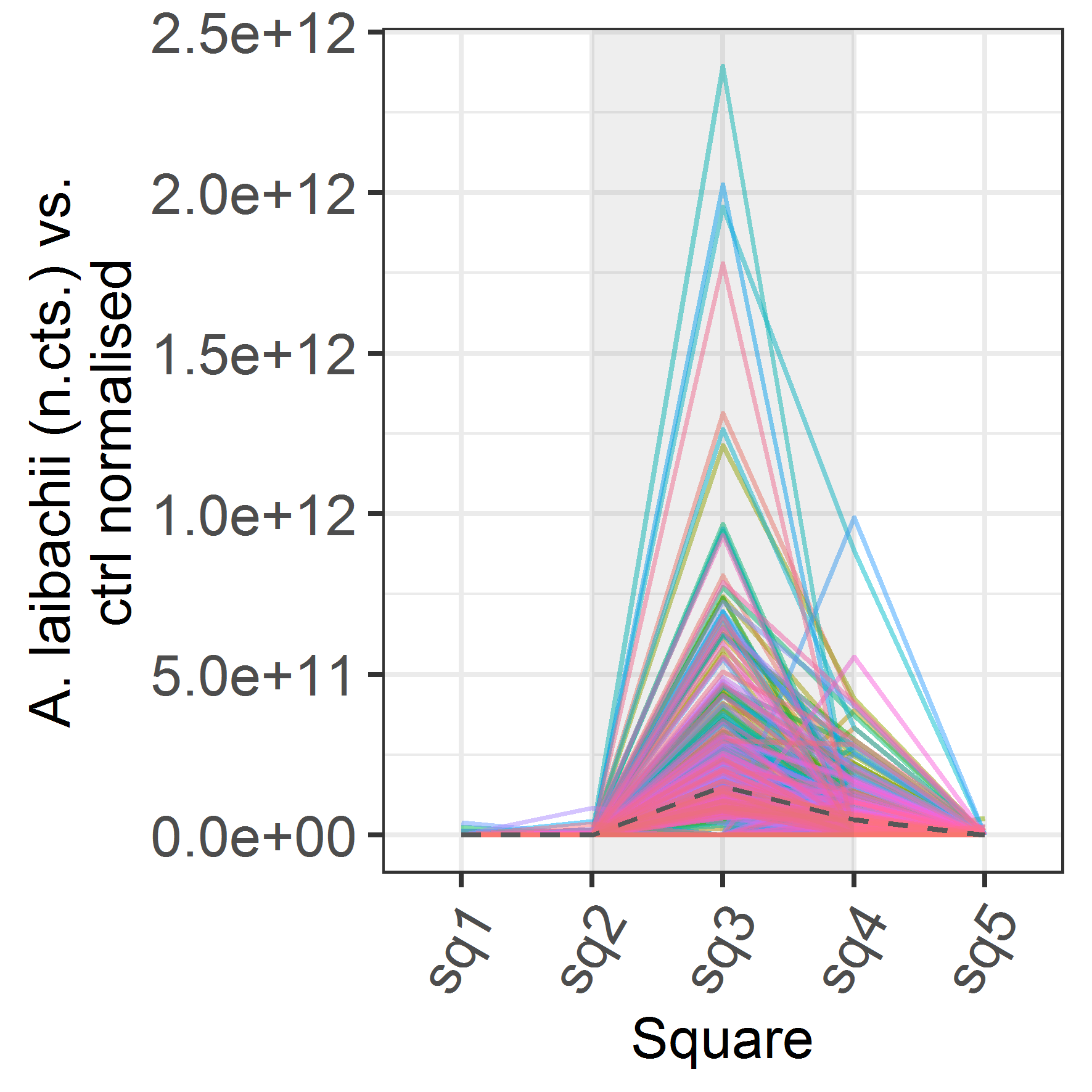

**Figure 6** Across leaf normalised expression plots of detected Albugo laibachii NC14 genes in replicate oom1. The grey dashed line shows the average expression of all detected 663 genes.

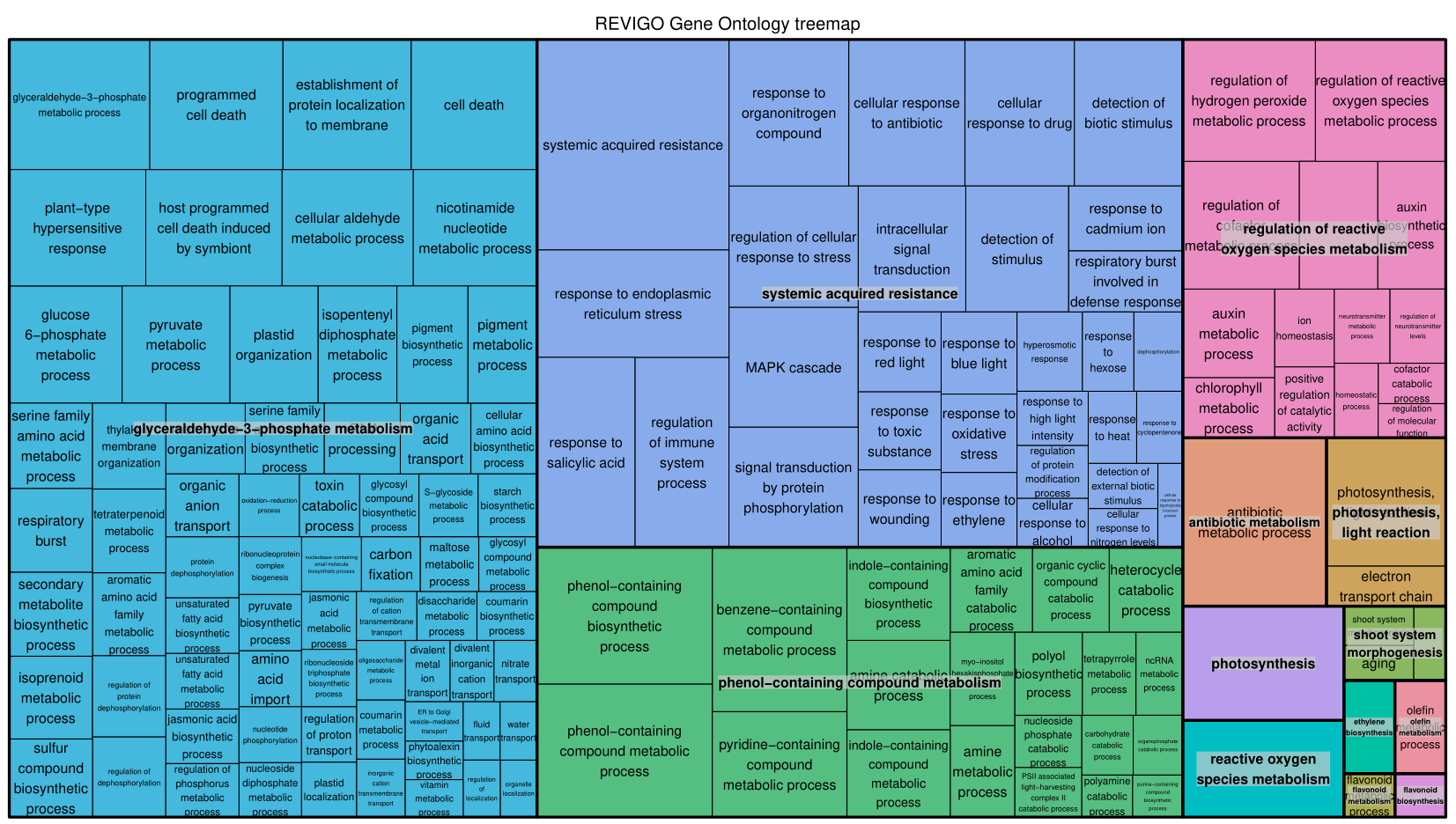

**Figure 7** Enriched GO-terms for the DE-genes of A. thaliana stressed with A. laibachii NC14 visualised with REVIGO [2].

### 1.6 Characterisation of spatial regulatory elements

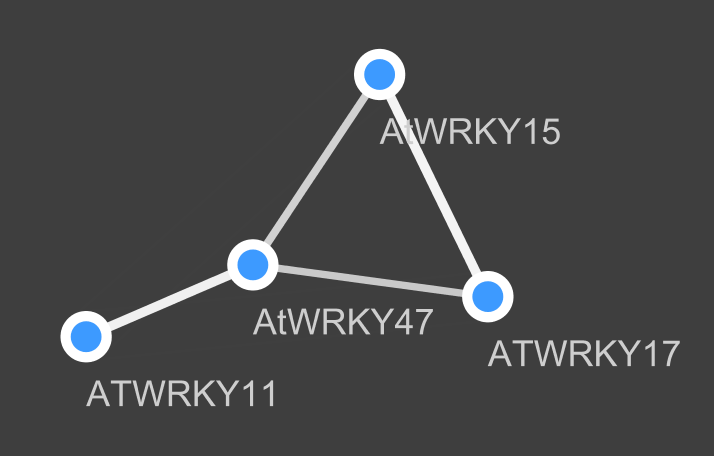

**Figure 8 WRKY Transcription factor network:** The transcription factors are linked to 388 of 526 total DE genes in the flg22 spotting experiment, the thickness of the line indicates the numbers of shared DE genes. WRKY11 and WRKY15 show spatially wider expression whereas WRKY47 and WRKY17 show spatially restricted expression.

### 1.7 *Grid-seq-1.1: Low Input RNA-seq (LORIS) library preparation*

### Grid-seq-1.1 workflow – mRNA extraction:

**Buffers:**

- Lysis buffer: (100mM Tris-HCl pH 7.5, 500mM LiCl, 10mM EDTA pH 8.0, 1% LiDS, 5mM DTT)
- Wash buffer A: (10mM Tris-HCl pH7.5, 0.15M LiCl, 1mM EDTA, 0.1% LiDS)
- Wash buffer B: (10mM Tris-HCl pH 7.5, 0.15M LiCl, 1mM EDTA)

Buffers are prepared with RNase free reagents.

(1) **Metal bead preparation for grinding**

- Wash beads with RNAse Zap
- Wash beads with DNAaway
- Wash beads multiple times with 80% EtOH
- Wash beads once with water
- Remove as much liquid as possible with a pipette tip
- Put beads in a sterile, screw-cap tube and dry on a 95 °C heating block
- Close tube and store beads until usage

(2) **Oligo-dT bead preparation for mRNA capture**

- Use 1µl Oligo-dT beads per reaction
- Wash beads twice in 200µl Lysis buffer (by vortexing well for washing and pelleting the beads on a magnet)
- Resuspend 1µl beads beads in 10µl Lysis buffer
- Add 10µl prepared Oligo-dT beads solution with lysis buffer to each well of a fresh 96-well plate (will be needed in step 3)

(3) **mRNA extraction procedure:**

Prepare the following grinding plate setup on dry ice.

- To each well with a sampled leaf section add 4-6x 1mm metal beads
- Dispense 10 µl lysis buffer to each well
- Close the wells with a round-cap PCR lid
- Transfer from dry ice immediately to the GenoGrinder
- Grind for 30 seconds with 1750 rpm on GenoGrinder
- Spin the plate for 10 seconds at 2000 x rcf and check if leaves were grinded successfully (a strongly green solution without solid material means successful grinding). If grinding was not successful, continue bead-bashing for another 10 seconds at 1750 rpm on the GenoGrinder.
- Spin the plate for 10 seconds at 2000 x rcf to remove air bubbles stemming from the lysis buffer and collect the liquid at the bottom each well
- Transfer lysed solutions into the plate with the washed mRNA beads but avoid carryover of metal ball-bearings.
- Close with the same round cap seals as used for grinding
- Shake the plate manually over-head to distribute buffer over the well
- Incubate on an over-head rotator for 10 minutes
- Collect beads on a 96-well magnet
- Wash twice with 50ul wash buffer A
- Wash once with 50ul wash buffer B
- Spin down briefly to collect the wash buffer at the bottom and remove the remaining wash buffer
- Elute in 8 µl water: incubate the sample at 80 °C for 2 minutes on a thermal cycler and immediately, move the plate back to the magnetic rack.
- Transfer the eluted mRNA to a fresh plate and store at -80 °C
- Use 2.5µl to proceed with the reverse transcription protocol

### Grid-seq-1.1 workflow – Reverse Transcription:

(1) **Oligo-dT binding**

Prepare the following mixture and add the dNTPs and the Oligo-dT30VN to the 2.5 µl mRNA.

|  | **Volume [µl]** |
| --- | --- |
| Oligo-dT30VN (10 µM) | 1.00 |
| dNTPs (10mM) | 1.00 |
| Purified mRNA | 2.50 |
| Total | 4.50 |

Incubate at 72 ⁰C in a thermal cycler for 3 min and snap cool on ice.

(2) **Reverse transcription**

Add the following to the Oligo-dT30VN bound mRNA:

|  | **Volume [µl]** |
| --- | --- |
| 5x FirstStrand buffer | 2.00 |
| Betaine (5M) | 2.00 |
| MgCl2 (1M) | 0.06 |
| Water | 0.09 |
| DTT (100mM) | 0.50 |
| RNAse inhibitor (40 U/µl) | 0.25 |
| TSO (100 µM) | 0.10 |
| SuperScript II (200 U/µl) | 0.50 |
| Total | 5.50 |

Run the following program: 42 ⁰C 90 min, (50 ⁰C 2 min, 42 ⁰C 2 min)15x, 70 ⁰C 15 min, 4 ⁰C forever.

(3) **PCR amplification**

Add the following to the reverse transcription reaction:

|  | **Volume [µl]** |
| --- | --- |
| Kapa HiFi HotStart Mix (2x) | 12.50 |
| IS PCR Primers (10 µM) | 0.25 |
| Water | 2.25 |
| Total | 15.0 |

Run the following program in a thermal cycler: 98 ⁰C 3 min, (98 ⁰C 20 s, 67 ⁰C 15 s, 72 ⁰C 6 min)15x-17x*, 72 ⁰C 5min, 4 ⁰C forever.

*This depends on the sample and the input amount. For 1 mm^2^ leaf sections 15 PCR cycles deliver high yields.

(4) **AMPure XP cleanup**

Perform a 0.64x AMPure XP cleanup (add 16ul AMPure XP beads to each reaction), elute in 20 ul 1x TE buffer (10 mM Tris-HCl, pH 8.0; 0.1 mM EDTA).

### Grid-seq-1.1 workflow – Nextera Reaction:

(1) **Prepration**

- Optional: Run a dilution series of Nextera enzyme with 1 ng DNA to estimate the appropriate amount of enzyme for tagmentation. Control yields on Qubit 2.0 HS and tagmentation sizes on the Bioanalyzer.
- Transfer 1 ng ds-cDNA per sample to a 96 well plate and dry the DNA in a drawer overnight (cover the 96-well plate loosely with the lid of a tip box).
- After drying and right before tagmentation prepare a fresh 96-well PCR plate containing the Illumina Nextera oligos with the appropriate barcodes (the concentration of the adapters in the subsequent PCR reaction should be 0.2 µM concentration):

|  | **Volume [µl]** |
| --- | --- |
| P5 forward (2.5 µM) | 2.00 |
| P7 reverse (2.5 µM) | 2.00 |

(2) **Tagmentation**

- Prepare the following mixture for tagmentation (on ice) and add immediately to the air-dried samples (on ice):

|  | **Volume [µl]** |
| --- | --- |
| Nextera enzyme | 0.10* |
| Illumina Nextera buffer (2x) | 2.50 |
| Water | 2.40 |
| Total | 5.00 |
| * or as determined by titration |  |

- Briefly vortex, immediately spin down and transfer for 5 min to a thermal cycler at 55 ⁰C.
- After the 5 min incubation step transfer the 5 µl of the tagmentation reactions to the primer plate.

(3) **Kapa 2G Robust amplification of tagmented product**

- Prepare the following mix and add to the tagmentation reaction + primer plate:

|  | **Volume [µl]** |
| --- | --- |
| Kapa 2G robust buffer (5x) | 5.00 |
| dNTPs (10 mM) | 0.50 |
| 2G robust polymerase (5 U/µl) | 0.10 |
| Water | 10.4 |
| Total | 16.0 |

Run the following program: 72 ⁰C 3 min, 95 ⁰C 1 min, (95 ⁰C 10 s, 65 ⁰C 30 s, 72 ⁰C 2 min 30 s)11x, 72 ⁰C 2 min 30 s, 4⁰C forever.

(4) **AMPure XP cleanup**

Perform a 0.64x AMPure XP cleanup (add 16ul AMPure XP beads to each reaction), elute in 20 µl 1x TE buffer (10 mM Tris-HCl, pH 8.0; 0.1 mM EDTA).

(5) **Library quantification and pooling**

Quantify the libraries using a Qubit2.0 Fluorometer or a device of similar sensitivity. Control randomly selected Illumina libraries on the Bioanalyser or a similar device able to visualize library profiles. Pool equimolarly before sequencing.
